# Construction and reconfiguration of dynamic DNA origami assemblies with coiled-coil patches and patterns

**DOI:** 10.1101/2023.09.23.559112

**Authors:** T. Teng, J. Bernal-Chanchavac, N. Stephanopoulos, C.E. Castro

## Abstract

DNA origami nanodevices achieve programmable structure and tunable mechanical and dynamic properties by leveraging the sequence specific interactions of nucleic acids. Previous advances have also established DNA origami as a useful building block to make well-defined micron-scale structures through hierarchical self-assembly, but these efforts have largely leveraged the structural features of DNA origami. The tunable dynamic and mechanical properties also provide an opportunity to make assemblies with adaptive structure and properties. Here we report the integration of DNA origami hinge nanodevices and coiled-coil peptides into hybrid reconfigurable assemblies. With the same dynamic device and peptide interaction, we make multiple higher order assemblies by organizing clusters of peptides (i.e. patches) or arranging single peptides (i.e. patterns) on the surfaces of DNA origami to control the relative orientation of devices. We use coiled-coil interactions to construct circular and linear assemblies whose structure and mechanical properties can be modulated with DNA-based actuation. Actuation of linear assemblies leads to micron scale motions and ∼2.5-10-fold increase in bending stiffness. Our results provide a foundation for stimulus responsive hybrid assemblies that can adapt their structure and properties in response to nucleic acid, peptide, protein, or other triggers.

## Introduction

Molecular self-assembly is a promising route to construct biomimetic or bioinspired materials that leverage the diverse properties and interactions of biomolecules (1). Nature produces many examples of self-assembled adaptive materials such as actin filaments that form bundles via local interactions with actin binding proteins, in order to modulate mechanical properties (2–4). Biomolecular nanotechnology provides a useful approach to mimic many of the structural, dynamic, and mechanical features of these systems. In particular, DNA nanodevices have been designed to exhibit programmed reconfigurations in response to a variety of triggers(5,6), for example to actuate the application of forces to biomolecules(7). Integrating these dynamic device functions into higher-order self-assembled architectures could provide an interesting approach to mimic the emergent properties of biomaterials such as adaptive structure and stiffness, large-scale shape changes, and stimulus-driven assembly or disassembly (8–10). Here we leverage the interaction properties of coiled-coil peptides and the structural and dynamic properties of DNA origami to make hybrid DNA-peptide constructs where reconfiguration of the DNA devices can regulate the structure and mechanical properties of higher order assemblies.

DNA has emerged as one of most common materials for the versatile self-assembly of precise nanostructures and dynamic nanodevices(11–14). In particular, DNA origami(15,16) leverages base-pairing interactions between a long ssDNA scaffold and hundreds of short ssDNA staples to create nanostructures with complex shapes and programmable motion (17,18). Here we take advantage of these features to construct higher order dynamic assemblies. Prior work has established the hierarchical assembly of DNA origami nanostructures to scale up the overall dimensions, demonstrating methods to make micron-sized assemblies with precisely organized components (19–21). Recent examples have further demonstrated building dynamic properties into these higher order assemblies; for example: controlling growth/disassembly(22); actuating changes in chirality(23), cross-section(24), bending(25), or length(26) in 1D assemblies; or changing shape in 2D assemblies(27,28). Building on these prior efforts, here we aimed to integrate actuation and assembly to make reconfigurable materials where actuation of DNA origami devices can control higher order assembly structure and mechanical properties.

Furthermore, as a step towards integrating the advantages of DNA-based assemblies with amino acid materials, we combined the structure and dynamic properties of DNA origami devices with the tunable interactions of peptides (29,30). Peptides have a number of useful properties such as biocompatibility(31); well-established approaches for sequence design(32) and synthesis (33), specific binding and self-assembly capabilities (34), and an existing basis for stimulus-responsive motifs(35). Here, we leverage the specific binding interactions of coiled-coil peptides. Coiled-coil interactions (36), which consist of binding between two α-helical peptides driven by hydrophobic and charge-charge interactions, and are one of the most abundant protein assembly motifs found in nature. Previous research has identified peptide sequences that lead to coiled-coil interactions with well-understood design principles(33). DNA-modified coiled-coils have been demonstrated as a good hybrid adhesive for building higher order structures, such as DNA origami filaments(37,38), as well as to integrate DNA nanostructures with functional proteins bearing complementary fused coils(39). Coiled-coils are also, in principle, reversible through an analogous mechanism to toehold-mediated strand displacement, through the addition of a fully complementary coil(40).

Expanding on this approach, here we employ coiled-coil interactions to construct assemblies that harness the reconfigurability of dynamic DNA origami devices. We demonstrate the ability to control the conformation of a dynamic DNA origami device with coiled-coil peptides. In addition, we used coiled-coil interactions as an adhesive to assemble multiple assembly configurations. To achieve these distinct configurations, we engineered specificity into the coiled-coil adhesion by patterning two peptides (which form a heterodimeric coiled coil) on the DNA origami surfaces to control the relative orientation of adjoining devices, enabling formation of distinct higher order assemblies from the same dynamic DNA origami building block (i.e. polymorphic assembly). Furthermore, we combined coiled-coil assembly with strand displacement actuation of dynamic DNA origami devices to modulate assembly structure and properties. Taken together, our results demonstrate hybrid DNA origami-peptide dynamic structures as useful building blocks for assemblies with polymorphic and reconfigurable structure and adaptive properties. Furthermore, the use of coiled-coils provides an entirely orthogonal interaction mode, which we found helped to mitigate aggregation that is a common challenge of higher order assembly with DNA sticky ends. In the future, this approach can also enable facile integration of functional proteins into the hierarchical nanostructures and allow these nanostructures to respond to protein cues to that modulate assembly, conformations, or properties.

## Results and discussion

We implement a previously developed dynamic DNA origami hinge design with tunable mechanical properties as the basis for our hybrid assemblies(41). The hinge arms are comprised of 20-helix bundles organized on an 8x3 square lattice cross-section with 4 internal helices removed from the middle layer (Figure 1A). In addition, there are ∼8-nm protrusions sticking out past the vertex on the back end of the two arms. The arms are connected by eight ssDNA linkers. Four of these connections are short 2 nucleotide (nt) linkers that form a rotation axis at the hinge vertex, while the other four are long 70 nt ssDNA linkers used to modulate the hinge properties. Two of the four long ssDNA linkers span across the inner layer near the vertex, and the other two long linkers span across the outer layer of helices at the back end of the protrusion. Previous work has established versatile approaches to modulate the hinge mechanical properties (i.e. the flexibility and minimum energy angle). For this work we implemented the version referred to as nDFS.A (41)(Supplemental Figure S1), which exhibits a flexible conformational distribution with most angles falling between 50-100° (Supplemental Figure S2).

**Figure 1.**
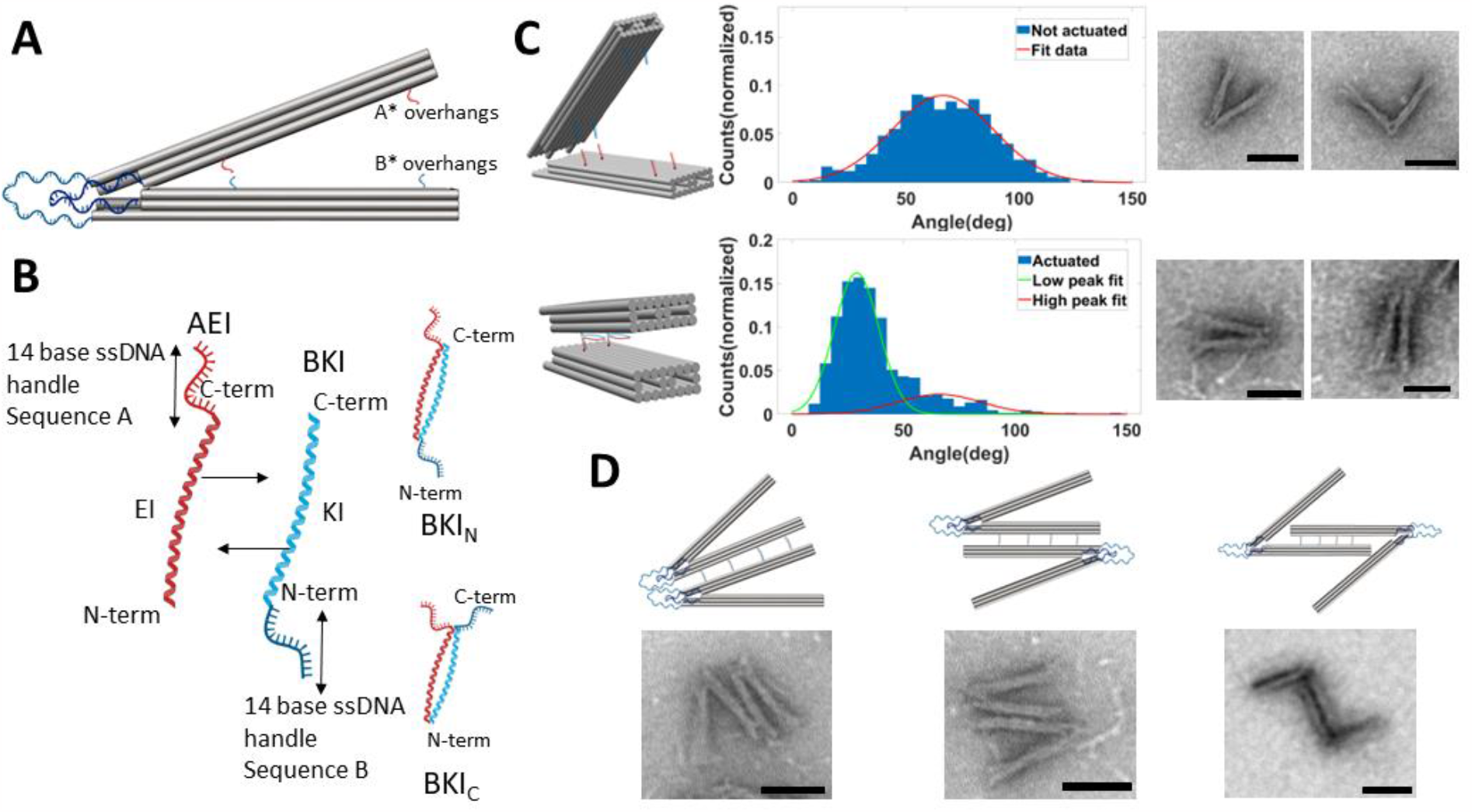
A) Schematic of the tunable DNA origami hinge design consisting of two arms, each a 3 × 8 helix square lattice bundle that is 210 bp long. The arms are connected by four flexible 70 nt long ssDNA linkers and four short 2 nt ssDNA linkers that define the axis of rotation. The arms can include two groups of ssDNA overhangs for binding coiled-coil peptides. B) Three DNA-peptide conjugates were used for actuation and assembly. The AEI conjugate has a 14 nt DNA handle ‘A’ attached to the coiled-coiled peptide EI on the C-terminal end. The other conjugate BKI has a 14 nt DNA handle ‘B’ attached to either the C-terminal (BKI_C_) or N-terminal (BKI_N_) of the KI coiled coil peptide. Combining the AEI and BKI conjugates yields a self-assembling coiled-coil interaction where the relative orientation of the handles depends on the use of BKI_C_ or BKI_N_. C) The angular conformations of non-actuated hinges, depicted schematically (top left), was quantified from TEM images (representative single hinges shown at right) revealing a flexible distribution of angles primarily in the range of 50-100° (Gaussian fit shown in red). Hinges actuated with coiled-coil interactions, shown schematically (bottom left), exhibited a large population of constrained angles in the range of 20-40°. Red and green lines show individual peaks of a double Gaussian model fit. D) Hinge dimers were assembled via AEI-BKI conjugates connecting the outer surfaces of the top and bottom hinge arms, leading to a mixture of dimers with hinges in the same orientation (left) or opposing orientations (middle). Locating the AEI-BKI interactions to the inner arm surfaces lead to dimers with only opposing orientations (right). Scale bars are 50nm.

We also implemented coiled-coil peptides that were previously utilized in the higher order assembly of static DNA origami nanostructures(37). These peptides consist of 28 amino acid residues consisting of four 7-residue (heptad) sequences; the peptide with heptad repeat EIAALEK is referred to as referred to as EI, and the peptide with heptad repeat KIAALKE is referred to as KI (Figure 1B). The ends of both EI and KI incorporate azidolysine (azK) residues for conjugation to ssDNA handles by strain-promoted azide–alkyne cycloaddition (SPAAC) chemistry. These two ssDNA handles, with orthogonal sequences termed as A and B, are 14-nt long and modified with dibenzocyclooctyne (DBCO) at the 5′ end through an amine linker. We followed established protocols(37) for the conjugation of ssDNA handle A to peptide EI (conjugate referred to as AEI) and conjugation of ssDNA handle B to peptide KI (conjugate referred to as BKI) (see methods for details). The handle A DNA oligo was conjugated on the C-terminus of the EI peptide, and handle B was conjugated on either the N-or C-terminus of the KI peptide leading to distinct binding configurations (Figure 1B) with different end-to-end distances between the DNA handles. The interaction where the DNA handles are on opposite ends (i.e. A on the C-terminus of EI and B on the N-terminus of KI), which we refer to as the N-terminal configuration, leads to a larger distance between the DNA handles, and the interaction where the handles are on the same end (i.e. A on the C-terminus of EI and B on C-terminus of KI) leads to a shorter distance between the DNA handles, which we refer to as the C-terminal configuration.

We aimed to establish the utility of peptide-peptide interactions for the actuation and assembly of dynamic DNA origami devices. To demonstrate actuation, we incorporated two sets of overhangs, one with sequences complementary to the A handle (A*) on the inner face of the top arm and another with sequences complementary to the B handle (B*) on the inner face of the bottom arm. We incorporated the overhangs for AEI and BKI attachment 38 bp away from the hinge vertex (Supplemental Figure S1). We chose to incorporate the C-terminal configuration of the coiled-coil interactions, which based on a rigid hinge model and length of conjugates would constrain the angle of the nanostructure to ∼35°. Therefore, binding of the coiled-coil peptides should provide a clear conformation change from the free hinge angle distribution (Figure 1C, top, Supplemental Figure S2).

Addition of the AEI and BKI_C_ DNA-peptide conjugates led to a large shift in the conformational distribution with most hinges exhibiting angles of ∼20-50°, as revealed by TEM image analysis (Figure 1C, bottom, Supplemental Figure S3), suggesting the coiled-coil interaction can effectively control the hinge conformation. We performed a mixed two-Gaussian model fit to the angle distribution data with one population exhibiting an angle of 29+-10° (i.e. average/peak ± standard deviation from Gaussian fit), corresponding to the actuated hinges, and a second population of hinges exhibiting an angle of 66+-19°, corresponding to hinges that remained unactuated. Based on these Gaussian fits we estimated an actuation efficiency of 74%. We also tested the coiled-coil actuation in the N-terminal binding configuration (AEI-BKI_N_). As expected, these results did not cause a major change in the angle distribution because the larger end-to-end distance of the ends of the coiled-coil attached to DNA would lead to a ∼70° angle. Nevertheless, a slight shift in the angle distribution still suggests some incorporation of the coiled-coil interaction (Supplemental Figure S4). In addition, we also tested designs where the overhangs for binding conjugates was positioned much farther from the hinge vertex (133 bp away) where we observed a larger population of hinges with angular conformations below 50° suggesting effective actuation (Supplemental Figure S5).

We then studied the use of coiled-coil interactions to make higher order assemblies of dynamic DNA hinge nanodevices. For higher order assemblies, we started with dimerization by making multiple sets of hinge devices each with one set of overhangs: one with A* overhangs on the outer face of the top arm, one with A* overhangs on the inner face of the top arm, one with B* overhangs on the outer face of the bottom arm, and one with B* overhangs on the inner face of the bottom arm (Supplemental Figure S6). We performed dimerization experiments by combining the outer A* and outer B* devices along with the AEI and BKI_C_ conjugates (we used BKI_C_ conjugates in all assembly experiments), which led to adhesion of two hinges in multiple possible configurations (Figure 1D, left and middle). Interestingly, we observed 66% of hinge dimers in the same orientation (i.e. vertices pointing in the same direction, Figure 1D, left, Supplemental Figure S7). Since the pattern of peptides on the surface is symmetric, either orientation should maximize EI and KI pairing. However, the same orientation dimer configuration might be favored due to slight twisting of the arms, steric interactions of the frayed ends, or possible weak base-pairing interactions of the ssDNA scaffold loops on the vertex end. We also designed a separate version of hinges where the coiled-coil interactions adjoin the inner faces of the arms on two separate hinges (Figure 1D, right, Supplemental Figure S7).

Expanding on the capability of coiled-coil peptides to assemble dynamic devices, we made new designs for the construction of higher order multi-device assemblies using only this specified coiled-coil peptide pair (AEI-BKI_C_) and a single hinge design. As a start, we first functionalized the outer side of one arm of the hinge with several A* overhangs and the other arm with B* overhangs, similar to the dimers, but in this case the AEI-BKI_C_ complexes could lead to self-assembly of larger multi-device assemblies. Figure 2A shows a representative TEM image (additional TEM images in Supplemental Figure S8) of the resulting assemblies, which showed having all AEI on the top arm and all BKI on the bottom arm led to a roughly even mix of neighboring devices that had similar orientations (i.e. vertex pointing in the same direction) or opposing orientations (i.e. vertex pointing toward opposing directions). We observed 51+-6% of assembly interactions led to neighboring devices in the same orientation. While we observed some circular (i.e. several devices in a row with the same orientation) and linear (i.e. several devices in a row with opposing orientations) assembly, these arrangements only persisted for a few devices, leading to irregular higher order structures. This mixed assembly results are likely due to the entire arm being coated with the same peptide, meaning the assembly interactions can randomly orient in either direction while still maximizing the pairing. While the assembly is efficient, there is no control over orientation, and hence no control over the higher order assembly structure.

**Figure 2.**
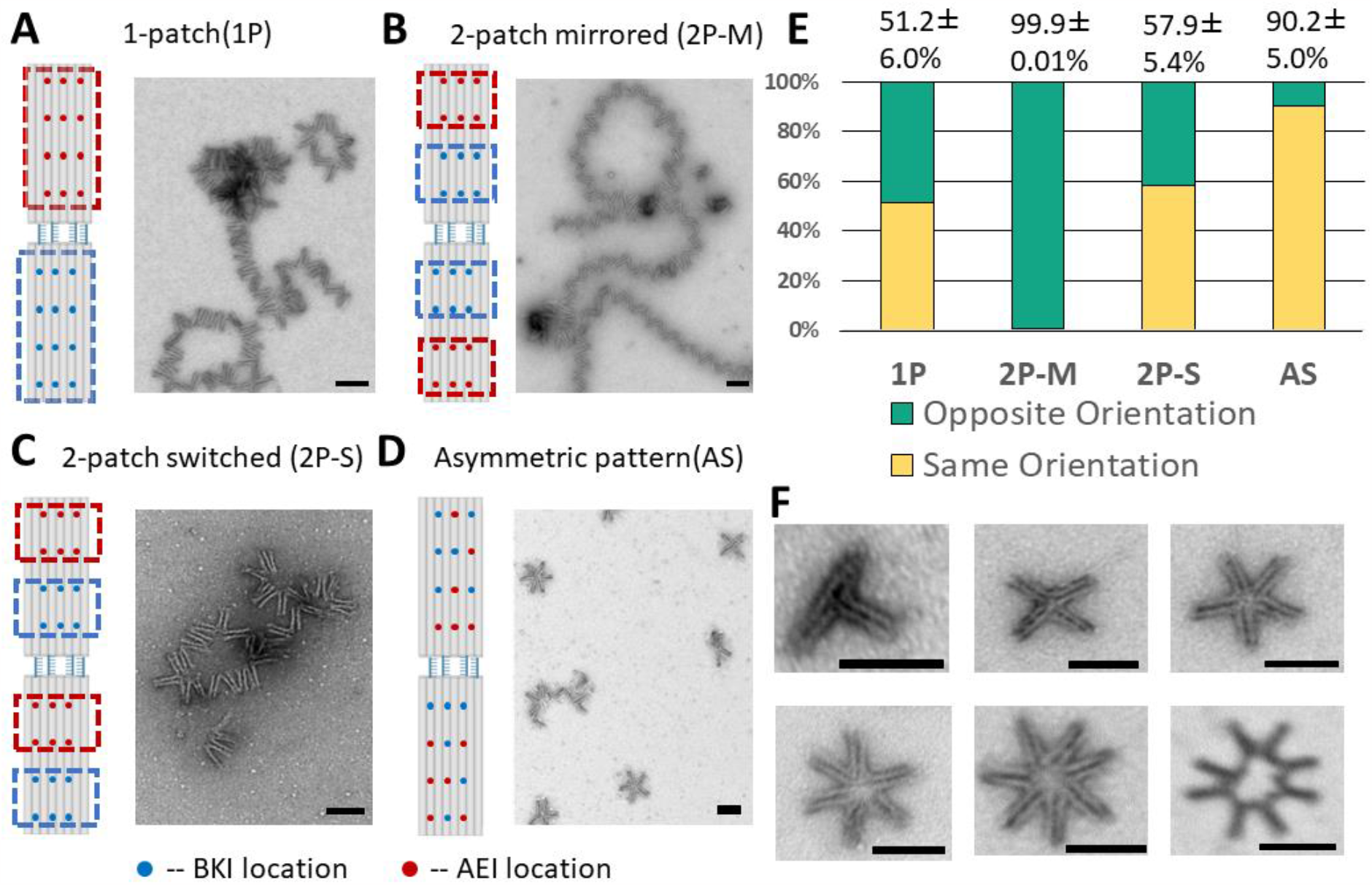
Controlling higher order self-assembly with peptide patches and patterns. A) Designing a patch of AEI peptides on the outer surface of the top arm and a patch of B* peptides on the outer surface of the bottom arm leads to higher order self-assembly. The hinges can attach in either orientation leading to assemblies with irregular structure as shown by TEM. B) Designing two patches per arm with BKI nearer the vertex and AEI farther from the vertex yields controlled assembly of devices in opposing orientations leading to long linear polymers as shown by TEM. C) Switching the arrangement of the patches on the bottom arm again leads to poor control over the orientation as shown by TEM, likely because either arm can attach to either arm. D) Engineering an asymmetric pattern of peptides facilitates assembly of hinges in similar orientation leading to circular assemblies as shown by TEM. E) The ratio fraction of neighboring devices that assemble in a similar (vertex pointing in same direction) or opposing (vertex pointing in opposite directions) was quantified from TEM images. F) Circular assemblies can form with different numbers of hinges from 3 up to 8 devices as shown by representative TEM images. Scale bars are 100nm.

To achieve uniform higher order assemblies, we engineered designs that could self-assemble into either controlled linear (i.e. all opposing orientation assembly interactions) or circular assemblies (i.e. all similar orientation assembly interactions). We took the approach of patterning A* and B* overhangs on the hinge arms to facilitate preferred binding in a specific orientation of neighboring hinges. We started by testing the two arrangements of overhangs shown in Figures 2B and 2C. Since the linear polymers require all opposing orientations, we arranged a simple pattern with a “patch” of B* overhangs nearer to the vertex and a patch of A* overhangs farther away from the vertex (both patches consist of 3 x 2 arrangement of overhangs) on the outer face of both arms. Hence, neighboring hinges would have to be in opposing orientations to form EI-KI interactions. TEM imaging revealed linear polymers were accurately formed with 99.9% of neighboring hinges exhibiting the correct opposing orientation (Figure 2B, Supplemental Figure S9) and long linear assemblies many over a micron in length, with the longest observed consisting of 78 hinge structures (Supplemental Figure S10). We also replaced the peptides to only DNA strands including same sticky ends in coiled-coil peptides to form self-assembled polymers, While this DNA sticky-end approach led to effective assembly, we observed more aggregation compared to the peptide assembly (Supplemental Figure S11). These results suggest the use of coiled-coil peptide interactions for higher order assembly can help overcome aggregation, which is a common challenge of forming large assemblies with purely DNA-sticky end interactions.

We tested two patch design approaches to facilitate a specific assembly of circular patterns. In the first approach, we reversed the pattern longitudinally in one arm of the hinge (i.e. the bottom arm was changed to have the A* patch near the vertex and B* patch away from the vertex). We found this case led to only 58+-5% of neighboring devices assembling in the same orientation, again leading to irregular higher order assemblies (Figure 2C, Supplemental Figure S12). This lack of specificity in assembly likely arises because the top arm can assemble with either the bottom arm of a different structure (leading to the same orientation) or the top arm of that structure (leading to an opposing orientation). To improve control over the same-orientation assembly, we reasoned that a more asymmetric pattern would avoid these undesired “top arm-top arm” and “bottom arm-bottom arm” assembly interactions, leading to higher efficiency of forming circular patterns. Towards this end, we tested the pattern shown in Figure 2D. TEM imaging revealed this complex pattern yielded drastically improved control over neighboring devices assembling with the same orientation, leading to efficient formation of circular patterns. Since the hinge is flexible (i.e. can adopt a range of angles), circular assemblies were observed containing a varying number of hinges. Figure 2F shows examples of circular assemblies containing different numbers of hinge devices (additional TEM images in Supplemental Figure S13). The circular assembly containing five hinges was the most abundant at 66%. This is consistent with the average angle of the unactuated hinge, which was about 70°.

We then tested our ability to control the number of hinges in circular assemblies by modulating the hinge angle. We introduced a closing DNA strand that actuates the hinge into a 41±13° angular conformation (Supplemental Figure S14), and we assembled these actuated hinges into circular patterns using the same asymmetric patterned self-assembly process. TEM images (Figure 3A, Supplemental Figure S15) revealed these constrained hinges led to an increase in the number of devices in the circular assemblies, with the majority of circles containing either 7x (37.9%) or 8x (32.0%) devices (Figure3B). Although the average hinge angle of 41° might suggest circular assemblies with 9 hinges would be likely, we only observed 7.4 % of circular assemblies containing 9 hinges. This suggests for smaller angles, the average angle does not completely govern the assembly process, likely due to the flexibility of the hinge and the thickness of the arms accounting for additional arc length subtended by each individual hinge. In addition, circular assemblies with a greater number of hinges have a higher probability of a defect (e.g. one opposing orientation assembly interaction), which will inhibit the formation of a fully closed circular assembly.

**Figure 3.**
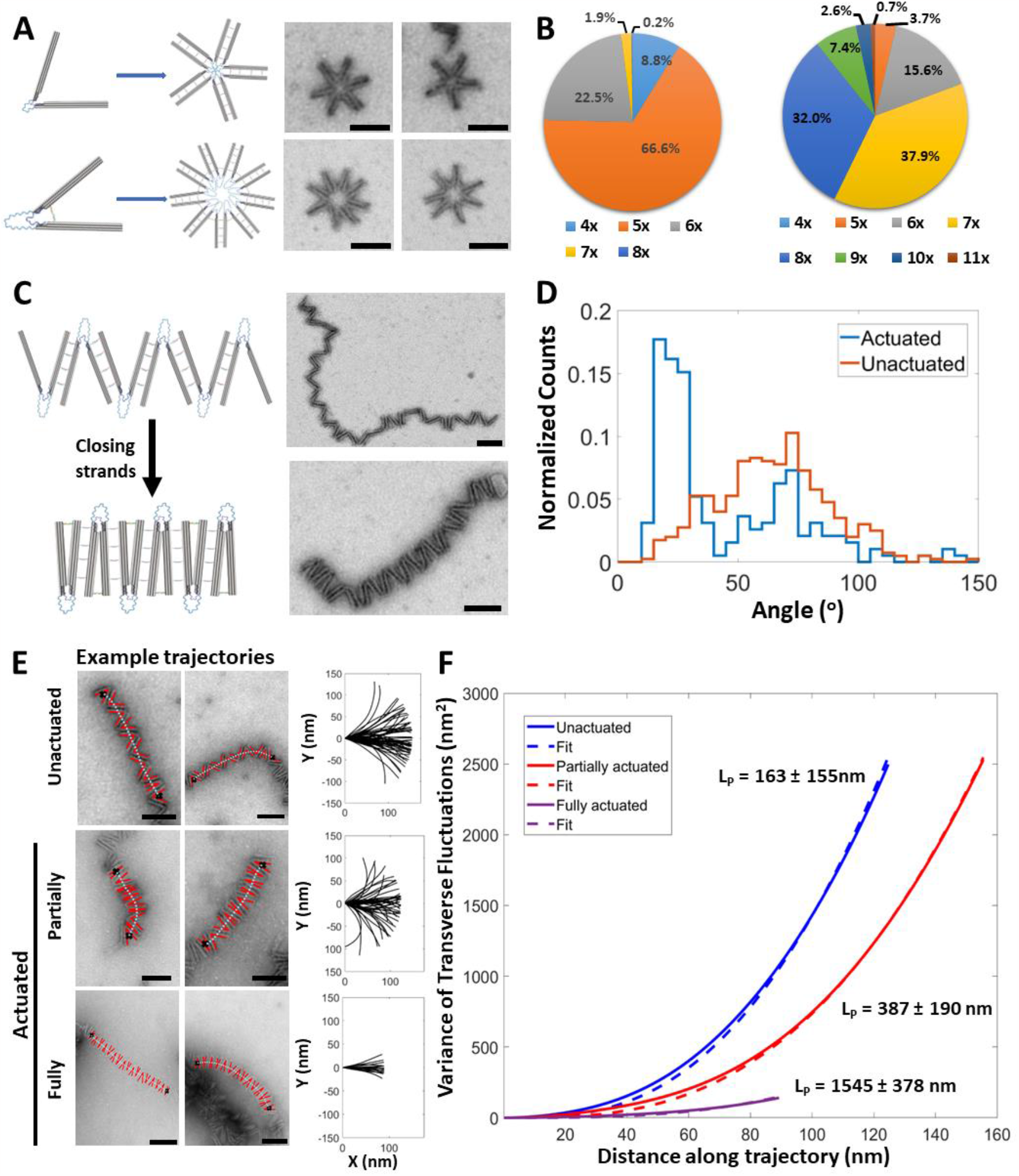
Characterization and control of higher order assemblies. A) Schematic and TEM images of circular assemblies made with free (top) or actuated (bottom) to adopt an angle of 41±13° using a DNA strand to form a strut. B) The actuated change to smaller hinge angles led to a shift from predominantly 5 hinges to predominantly 7 or 8 hinges in the circular assemblies. C) Schematic and TEM images of unactuated and actuated linear assemblies where hinges contained DNA overhangs on the inner faces of the arms and DNA closing strands were added after assembling to actuate the polymers. TEM images reveal efficient actuation with actuated regions appearing straighter than the unactuated polymers. D) TEM images analysis enables quantification of the variation of hinge angle distributions between non-actuated and actuated polymers. The actuation leads to a population of hinges with an angle of 20+-2 ° from 62+-5°. E) Examples of trajectory analysis of hinge polymers to evaluate the effects of actuation on the effective persistence length of polymers. The top row is the unactuated hinge polymer. The second row is partially actuated hinge polymer, which means in the trajectories 10 hinges were picked as same as in unactuated ones. The third line is fully actuated hinge polymer, which means in the trajectories only actuated hinges were picked no matter how many hinges assembled together. F) Analysis of hinge polymer trajectories revealed that the unactuated polymers exhibit an effective persistence length of 163±155nm. Partially actuated hinge polymers exhibited a persistence length of 387±190nm, and fully actuated hinge exhibit a persistence length of 1545±378nm, showing that the actuation leads to stiffer filaments. Scale bars are 100nm.

With these circular assemblies, we showed that *a priori* actuation to modulate the hinge structure can tune the structure of the higher order assembly. We next used the linear polymers to demonstrate dynamic control over the structure and properties of the fully formed assemblies. We created linear polymers with hinge devices containing DNA overhangs on inner facing sides of the hinge arms, which allow for closing of the hinge by introducing an additional DNA strand that latches the two arms together by base-pairing overhangs on each arm (Figure 3C, top). Experiments on isolated hinges showed that this actuation led to hinges with angular conformations of about 15+-4° (Supplemental Figure S16). We observed a similar distribution of angles for hinges incorporated in these linear assemblies compared to isolated unactuated hinges and we found that actuating linear assemblies led to the angle distribution of hinges in the polymers shifting from 62+-5° for the free hinges to 20+-2° for the actuated hinges (Figure 3D).

In addition to modulating the structure (i.e. changing end to end distance), the actuation led to assemblies that exhibited relatively less curvature (Figure 3C, Supplemental Figure S17), indicating an increase in stiffness. To quantify this increase in stiffness, we measured an effective polymer persistence length for the linear assemblies with or without actuation. We manually identified and traced the hinges along the polymer, and fit a spline curve to represent the trajectory along the center of the hinge (Figure 3E and Supplemental Figure S18). We calculated the variance of the transverse fluctuations from these trajectories, which can be correlated to the persistence length as previously described by Isambert et al (42). Fitting the transverse fluctuations of these trajectories resulted in a persistence length of 163±155 nm for the free hinge assemblies (Figure 3F). For the case where we added actuation strands, we observed incomplete actuation. The angle distribution (Figure 3D) for the hinges in the actuated polymers suggests an actuation efficiency of 72% (using a cutoff of 30°). The analysis of transverse fluctuations of these trajectories revealed a persistence length of 387±190 nm for the assemblies when we included hinges that remained open in the trajectory analysis (i.e. partially actuated hinge polymers, Figure 3F-E). While these results show a clear increase in stiffness, we hypothesized that the unactuated hinges had a significant impact on the assembly properties. To quantify the properties of fully actuated regions, we traced sections of polymers where all hinges were actuated, which resulted in a persistence length of 1545±378 nm (i.e. full actuated, Figure 3F-E). We interpret this result as indicative of the stiffness over a short distance where hinges are fully actuated, which likely also indicates the limiting behavior if the efficiency were optimized to 100% hinge actuation. To further explore these results, we simulated hinge polymer trajectories using the geometry of the hinges and experimentally measured angle distributions for the free hinge, partially actuated hinge (i.e. full angle distribution for actuated polymers in Figure 3E), and fully actuated hinge (i.e. only selecting angles below 30o from the angle distribution for actuated polymers in Figure 3E). Our simulation results (Supplemental Figure S18) yielded persistence lengths of 95nm, 284nm, and 5141nm for the free, partially actuated, and fully actuated polymers. These results are consistent with our experimental results providing further support for the effects of actuation and actuation efficiency on the mechanical properties. Overall, these results illustrate that these dynamic DNA origami-peptide hybrid assemblies can be actuated to modulate both structure and mechanical properties.

## Conclusion

In this work, we established the use of coiled-coil peptide interactions in controlling higher-order assembly of DNA nanodevice conformations, as well as constructing reconfigurable DNA assemblies with dynamic control over structure and mechanical properties. We used the coiled coil peptides as an adhesive where specificity of interactions can be controlled by the location and geometry of peptide interactions. We took advantage of the site-specific addressability of DNA origami structures to organize individual, patches, or patterns of peptide interactions to direct coiled-coil interactions for actuation or assembly. This result in particular highlights that even with a single peptide-peptide interaction (consisting of two complementary coils) we could use spatial organization on the origami to drive a range of specific multivalent interactions. We demonstrated isolated coiled-coil interactions can actuate device conformation changes, and patches or patterns of peptides can enable controlled higher order assemblies. We formed linear assemblies of hinge devices with simple arrangement of peptide patches, while circular assemblies required a more complex pattern of peptides to avoid spurious interactions. Furthermore, our results suggest using coiled-coil interactions for higher order DNA origami materials assembly can mitigate non-specific aggregation, which is a common challenge when using DNA sticky ends for higher order assembly. Finally, we showed the structure and mechanical properties of these assemblies can be modulated by controlling the dynamic device conformation either before (for circular assemblies) or after (for linear assemblies) the higher order self-assembly. We showed a 2-3 fold increase in linear polymer bending stiffness with actuation, but our results indicate that further optimization could lead to 10-fold or more increases in stiffness with actuation.

Our results build on recent efforts to make controllable assemblies of nanostructures(43,44) . Many studies have demonstrated higher order assembly of static DNA origami structures(19,20,45), and recent studies have extended to dynamic assemblies(22–27) . Inspired by the development of peptide-DNA conjugates for DNA origami assembly (39), here we used coiled-coil peptides as a useful adhesive for higher order self-assembly of dynamic devices where orientational control, and hence assembly structure control, can be engineered by organizing peptides into patches or patterns. Recent interest and advances in the colloidal or patchy particle assembly of DNA origami structures(46,47) can provide further guidance towards more complex and functional assemblies. In addition, we combined coiled-coil assembly with DNA strand displacement actuation in sequence, either “actuation-then-assembly” to modulate the structure of self-limiting circular assemblies, or “assembly-then-actuation” to modulate the structure and mechanical properties of linear assemblies. Prior work has demonstrated DNA origami assemblies with variable mechanical properties(48,49); however, these were not actuated systems, but rather different folded versions of structures that combined into higher order assemblies. The responsive nature of our assemblies and the combination of peptide and nucleic acid interactions to achieve assembly and actuation suggest these dynamic assemblies could serve as a useful platform to design adaptive materials that change shape or properties in response to amino acid or nucleic acid-based triggers or other environment stimuli. In particular, the use of coiled-coils could be an attractive approach for interfacing with a variety of proteins (39).

## Acknowledgements

This research was primarily supported by funding from the Department of Energy through grant number DE-SC0017270 to C.E.C. Research reported was also supported by The National Institute of General Medical Sciences of the National Institutes of Health under grant number DP2GM132931 to N.S. The content is solely the responsibility of the authors and does not necessarily represent the official views of the Department of Energy or the National Institutes of Health. Images shown were generated at the Campus Microscopy and Imaging Facility at The Ohio State University. This facility is supported in part by grant number P30 CA016058, National Cancer Institute, Bethesda, MD. We would also like to thank members of the Castro and Stephanopoulos laboratories for valuable feedback.

