## Supplemental for "Construction and reconfiguration of dynamic DNA origami assemblies with coiled-coil patches and patterns"

### Materials and Methods

### DNA origami folding and purification

The DNA origami hinge was designed as combination of an 8064 nt ssDNA scaffold(1)and staple strands by using software caDNAno(2). The design diagram from caDNAno is included in Supplementary Figure S1, illustrating all design details and the sequences for all staples are included in Table S1. Briefly, the parts of overhangs used for binding coiled-coil peptides conjugates are fully complementary to the DNA handles in conjugates. For AEI peptides, the overhang A* sequence is GTAATACCAGATGG and for BKI peptides, the overhang B* sequence is TATATGGTCAACTG. In addition, overhangs on the inner side of arms in self-assembly design are used for binding close strands to actuate the structures. The closing strand sequence is AGTGGACCAGTGGGTCTTCGTATAGACCCGACTTTGGGCCTAAGTGGGTCCACACGCACG. The scaffold was prepared in our laboratory[ref], and the staple strands were ordered from a commercial vendor (IDT, Coralville).

Folding reactions contained 20 nM scaffold and 200 nM of each staple strand in a ddH2O solution containing 1× FOB and 20 mM MgCl2.This folding reaction was subjected to thermal annealing in a thermal cycler (Bio-Rad, Hercules, CA) consisting of rapidly heating the solution to 70°C for 15 min, followed by annealing over the range of 63–57°C for 3 hours per degree Celsius, and then cooling for 30 min at 4°C.

The structures were purified by centrifugation in a polyethylene glycol (PEG) solution (3). The production of folding reaction with folded DNA origami structures was mixed by an equal volume of PEG buffer (15% PEG MW8000, 200 mM NaCl and 100 mM Tris). The mixture was then centrifuged at 4 °C and 16000g for 30 min. The supernatant was removed, and the origami structures were resuspended in 1× FOB with 20 mM MgCl2. Then the concentration of the structure in the solution was measured by Nanodrop (NanoDrop 2000C Spectrophotometer, Thermo Scientific) in 260 nm absorbance.

**Actuation**

Purified individual structures with group 1 or group 2 overhangs were mixed with 20× excess AEI and BKI conjugates relative to the structure concentrations, and then the final concentration of structures was adjusted to 5 nM with buffer containing 1× FOB and 20 mM MgCl2 and final volume is 50 µl. The mixture was subjected to an incubation of 15 hours at 37 $^{o}$C.

**TEM and analysis**

The sample was diluted to 1 nM in 0.5× TBE with 10 mM MgCl2 buffer for transmission electron microscopy (TEM). For TEM grid preparation, 4 µl of sample volume was deposited on Formvar-coated copper TEM grids, stabilized with evaporated carbon film (Ted Pella; Redding, CA). Then the sample was incubated on the grid for 4 min when incubating single structure, and for 8 min when incubating self-assembly structures, and then was wicked away with filter paper. The sample was then stained with 2% uranyl formate (SPI, West Chester, PA, USA). First, a 10 µL drop was applied for 2 s and wicked away to wash the sample, and then another 10 µL drop was applied for 15 s and then wicked away with filter paper. TEM imaging was performed at the OSU Campus Microscopy and Imaging Facility on an FEI Tecnai G2 Spirit TEM using an acceleration voltage of 80 kV at different magnifications.

The raw TEM figures were organized into a gallery containing clear formations (Figure S2-S6,S14-S16) by using the particle picking tool in software EMAN2. Then the angles were measured in the software ImageJ by drawing two straight lines directly on each particle along the inner side of each arm in one hinge.

We used MATLAB as the postprocessing tool to convert the angle data sets to probability density histograms. To estimate the fraction of the hinge in non-actuated or actuated states, we used the peakfit Matlab function program(4), which uses a non-linear optimization algorithm to decompose a complex, overlapping-peak signal into its component parts, by assuming the conformational distribution consisting of two populations. The peak number was set as 2 for the actuated sample and the distribution was fit with a combination of two Gaussian distribution, termed as the actuated and non-actuated parts.

**Self-assembly**

Purified individual structures with each designed overhang arrangement were mixed with 20× excess AEI and BKI conjugates relative to the structure concentrations, and then the final concentration of structures was adjusted to 5 nM with buffer containing 1× FOB and 20 mM MgCl2 and final volume is 50 µl. For the linear self-assembly polymer, the mixtures were then subjected to a 1-cycle low temperature annealing ramp starting at 45 °C followed by an anneal phase at −2 hours per °C until 20 °C, to prevent extensive aggregation. For other self-assembly structures, the process was 2-cycle annealing ramp, which was repeated twice.

**Circle pattern actuation**

Before self-assembly process, the purified structures were mixed with 0.5 µl of 10µM closing strands and then incubated at 37 $^{o}$C for 15 hours to yield an angle of hinges mostly actuated to 45$^{o}$.

Linear polymer actuation

After self-assembly process, 0.5 µl of 10µM closing strands were added to the production. And then the mixture was incubated to 37 $^{0}$C for 15 hours.

**Persistence length from shape variance in TEM images**

TEM images of non-actuated and actuated self-assembly polymers were analyzed using MATLAB. To discretize the shape of the polymers, 11 points on the connection of the hinge along the trajectory were manually selected by clicking on the image Figure S17A and Figure S18A These selected points were used to fit a cubic spline of the trajectory coordinates along the filament path to obtain fine resolution of the curvature. Configurational distributions were obtained by aligning the filament trajectories so that they started at the origin and initially pointed in the horizontal direction Figure S17B and Figure S18B. Isambert et al. derived a relation between the filament persistence length (LP) and the average transverse fluctuations, $\langle [D(s)]^2 \rangle$,

$$\langle [D(s)]^2 \rangle = L_p^2 \left[ 2\frac{s}{L_p} + \frac{16}{3}\exp\left(-\frac{s}{2L_p}\right) - \frac{1}{3}\exp\left(-\frac{2s}{L_p}\right) - 5 \right] \quad (1)$$

The average transverse fluctuations were determined as a function of arc length from the filament configurational distributions. The LP of the actuated and non-actuated self-assembly polymer were characterized respectively by calculating the average of the transverse fluctuations squared from the configurational distributions and fitting Eq. (1). Figure S17 shows the model of non-actuated polymers fits compared to the data, which resulted in LP of 327 nm. Figure S18 shows the model of actuated polymers fits compared to the data, which resulted in LP of 536 nm.

**<u>Materials and supplies</u>:** Fmoc-protected amino acids for peptide synthesis were purchased from EDM Millipore. Fmoc-azidolysine was purchased from Combi-Blocks Inc. Dichloromethane (DCM) was purchased from Millipore Sigma. Dimethylformamide (DMF) and diethyl ether were purchased from Oakwood Chemical Inc. Piperidine was purchased from Alfa Aesar. Oxyma and DIC were purchased from ChemImpex. TFA was purchased from Oakwood, and Rink Amide resin was purchased from Novabiochem. DBCO-sulfo-NHS linker was purchased from Click Chemistry Tools. All oligonucleotides were purchased from Integrated DNA Technologies.

**<u>Peptide synthesis and characterization</u>:** Peptides were obtained using solid phase peptide synthesis (SPPS) on a CEM Liberty Blue instrument. Synthesis was performed on a solid phase Rink-Amide resin (0.78 mmol/g) at a 0.1 mmol scale, using a standard Fmoc protocol and deprotected in 20% piperidine in DMF. Amino acids, coupling agents, DIC and Oxyma, were added in a 10-fold molar excess. Crude peptides were cleaved by shaking the resin by in a solution containing trifluoroacetic acid (TFA), triisopropylsilane (TIPS), and water in a ratio of 95:2.5:2.5 for 3.5 h. The resin was washed with TFA and concentrated under nitrogen. The solution was then added to 40 mL of cold diethyl ether to precipitate the peptide. The solution was centrifuged at 4200 rpm for 10 min, the supernatant was removed, and the pellet was allowed to dry overnight. The dried pellet was dissolved in a mixture of water and acetonitrile (50:50) and 0.1% TFA. Peptides were purified via reverse phase chromatography on a Waters HPLC using a Phenomenex column with C18 resin. A linear gradient was generated using water/acetonitrile + 0.1% TFA, from 10% to 100% acetonitrile over 50 minutes. Peak fractions were collected based upon their absorbance at 230 nm and tested for purity by MALDI-TOF mass spectrometry on a Bruker Microflex LRF MALDI using α-cyano-4-hydroxycinnamic acid matrix (Sigma). Pure fractions were pooled and lyophilized, and peptides were stored at -20 °C until use.

**Synthesis of Peptide-DNA Conjugates** DNA-peptide conjugates were prepared via strain-promoted azide-alkyne cycloaddition (SPAAC) following reported protocols (5). Briefly, amine modified oligonucleotides were dissolved in phosphate buffered saline (PBS) to a concentration of 1 mM. A 10 molar excess of DBCO-sulfo-NHS dissolved in DMSO was then added to the DNA and agitated at RT overnight. The reaction mixture was washed six times with a 3 kDa molecular weight cutoff (MWCO) filter (Amicon) to remove any excess DBCO. The resulting DNA was purified by reverse phase HPLC on an Agilent 1220 Infinity LC HPLC with a Zorbax Eclipse XDBC18 column. A Linear gradient was generated using 50 mM TEAA/Methanol from 10% to 100% methanol over 60 minutes. Peak fractions were collected based upon their absorbance at 260 nm and tested for purity by MALDI-TOF mass spectrometry on a Bruker Microflex LRF MALDI in 3-Hydropicolinic acid (HPA) matrix (Sigma). Pure fractions were pooled, and buffer exchanged into water using a 3 kDa MWCO filter. The purified DBCO-DNA was then mixed with the azido-peptides, heated to 37 °C, and shaken overnight. Following the reaction, samples were exchanged into water using a 3 kDa MWCO filter, and purified by reverse phase HPLC, similar to the modified DBCO-DNA.

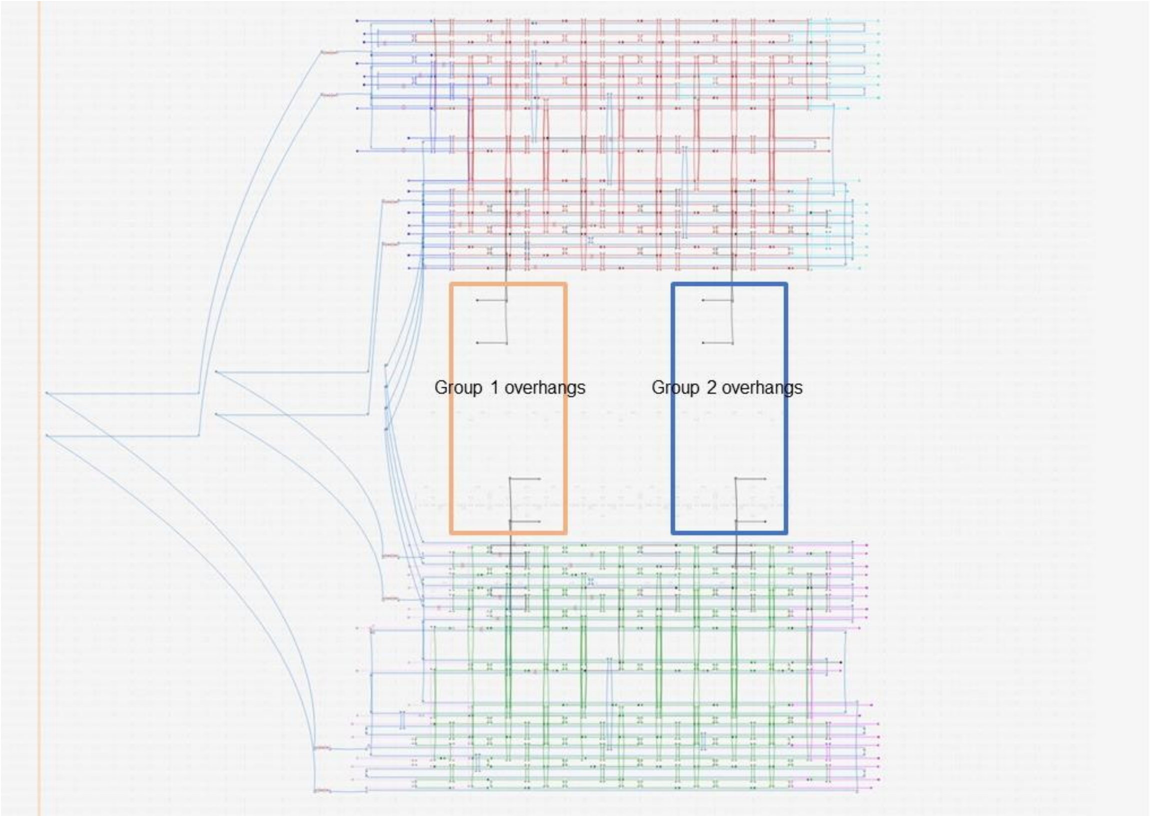

**Figure S1.** The caDNAno schematics of DNA origami hinge. The labeled areas are group 1 and 2 overhangs for binding AEI and BKI peptides to actuate the structure.

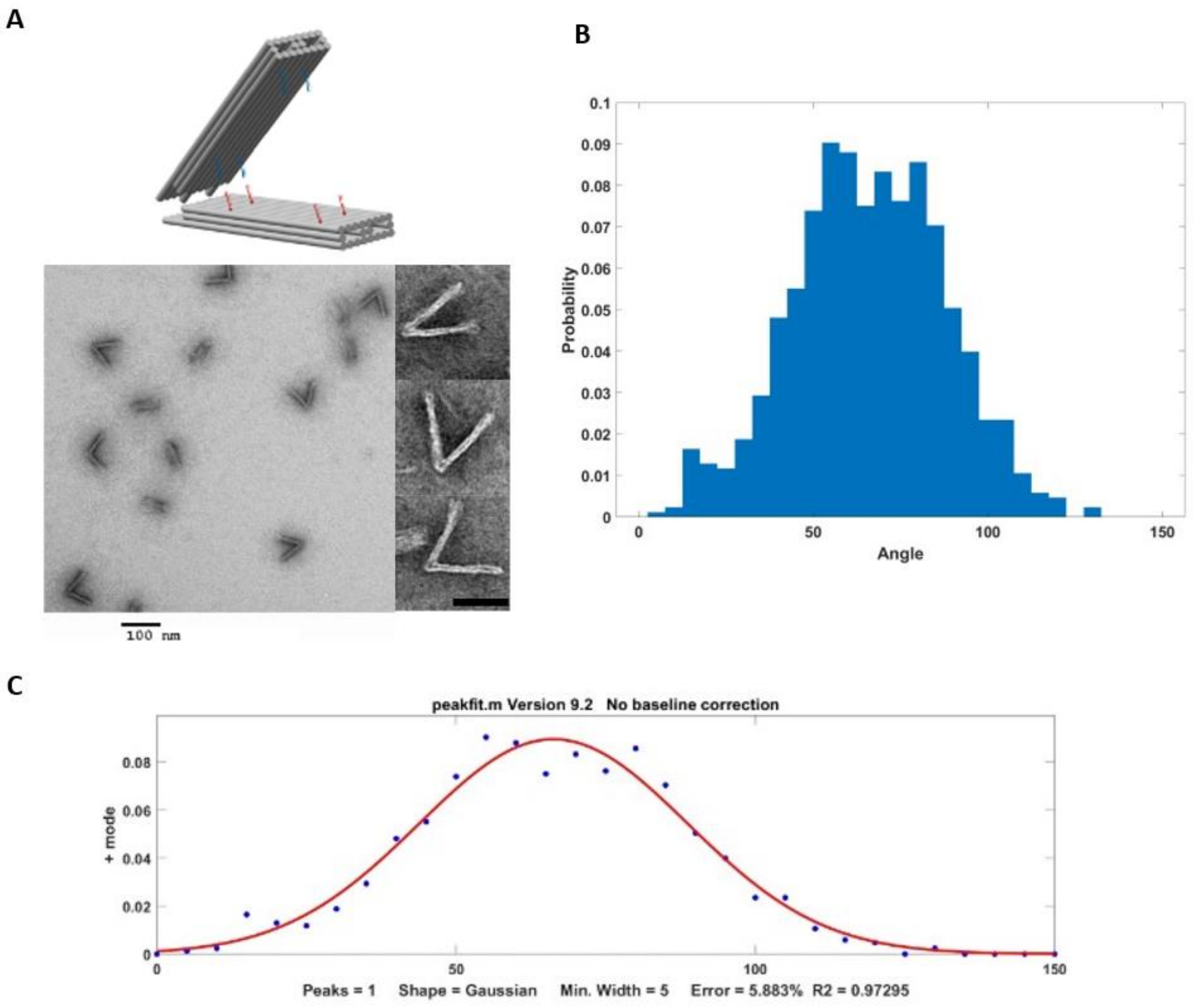

| Position | Std | Area |
|---|---|---|
| 66.2 | 22.4 | 5.0 |

**Figure S2.** TEM figures and analysis of non-actuated DNA origami hinge. A) Examples of TEM figures. B) The angular distributions obtained from TEM results. C) Single peak fitting of the angular distribution by using Matlab function program.

A

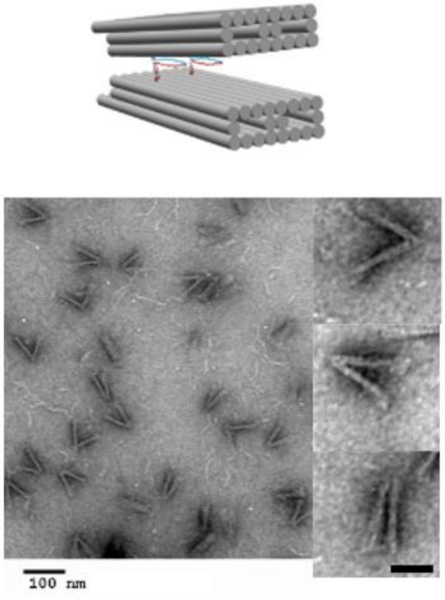

B

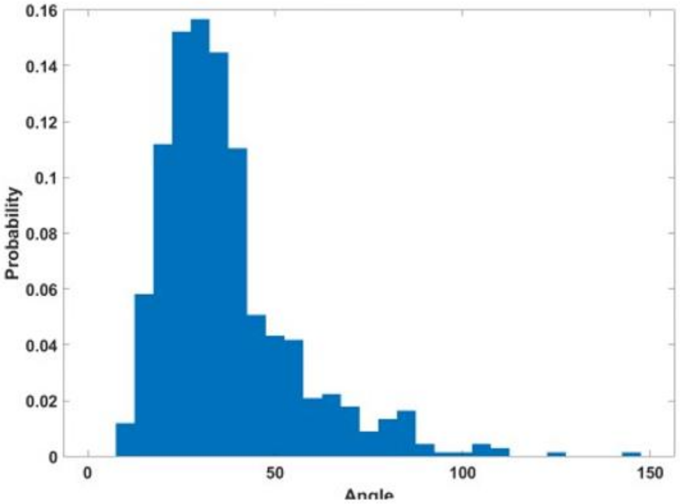

C

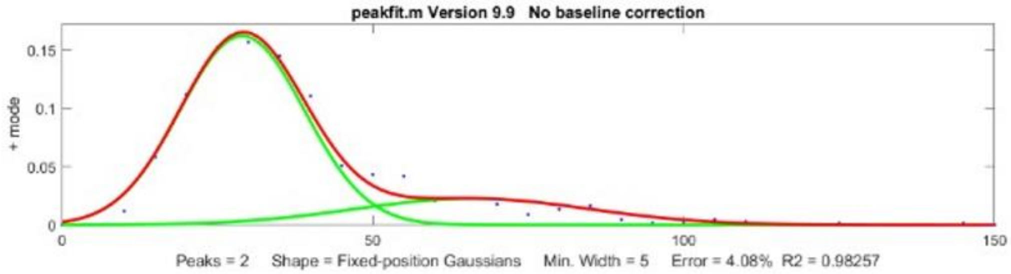

| | Position | Std | Area |
|---|---|---|---|
| High peak | 66 | 18.8 | 1.3 |
| Low peak | 29 | 10.0 | 3.6 |

**Figure S3.** TEM figures and analysis of actuated DNA origami hinge when C-term coiled-coil peptides binding group 1 overhangs. A) Examples of TEM figures. B) The angular distributions obtained from TEM results. C) Double peaks fitting of the angular distribution. The lower peak is $29.0^{o}$ and the higher peak is $54.11^{o}$.

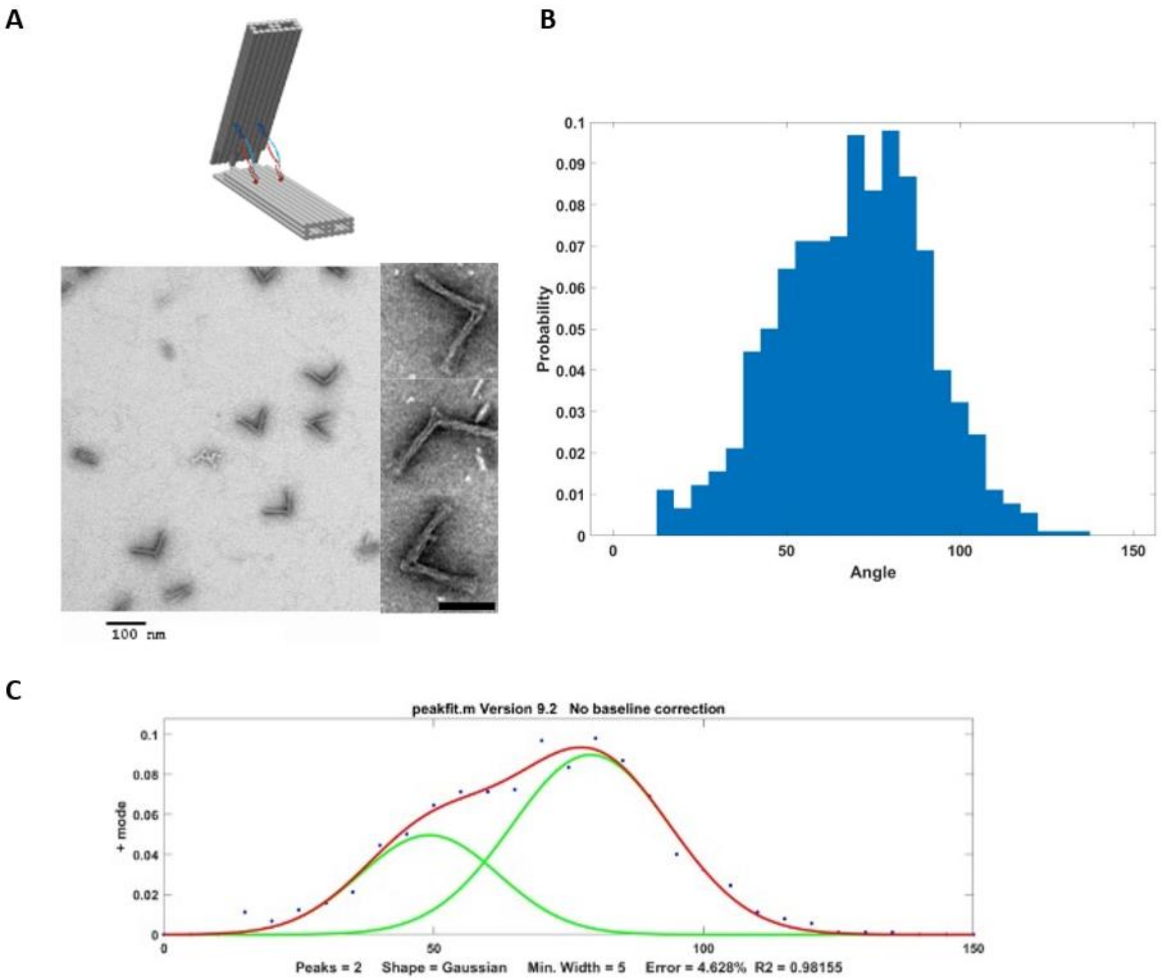

| | Position | Std | Area |
|---|---|---|---|
| High peak | 79.1 | 14.6 | 3.3 |
| Low peak | 49.1 | 12.8 | 1.5 |

**Figure S4.** TEM figures and analysis of actuated DNA origami hinge when N-term coiled-coil peptides binding group 1 overhangs. A) Examples of TEM figures. B) The angular distributions obtained from TEM results. C) Double peaks fit of the angular distribution. The lower peak is $49.2^{o}$ and the higher peak is $79.1^{o}$.

A

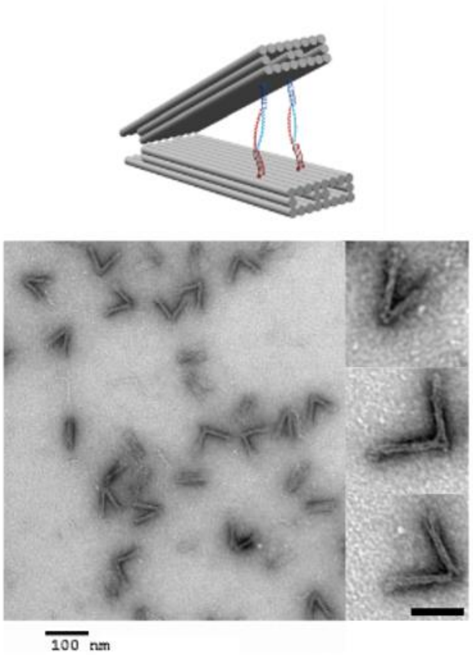

B

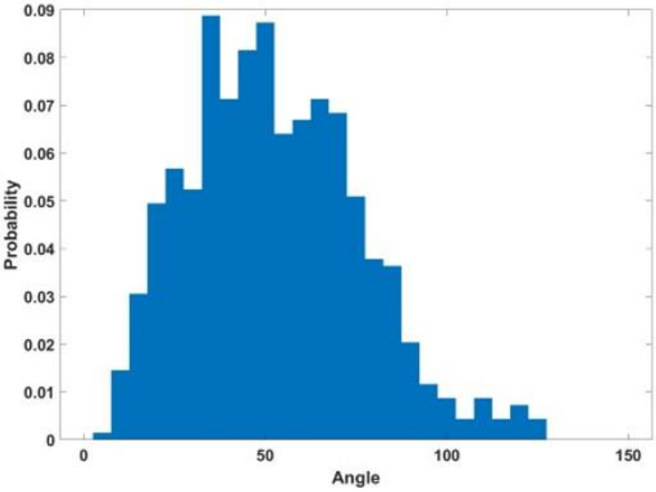

C

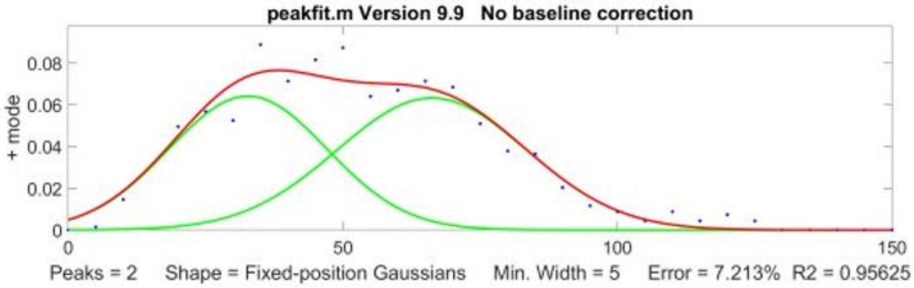

| | Position | Std | Area |
|---|---|---|---|
| High peak | 66.2 | 17.2 | 2.7 |
| Low peak | 32.7 | 14.4 | 2.3 |

**Figure S5.** TEM figures and analysis of actuated DNA origami hinge when N-term coiled-coil peptides binding group 2 overhangs. A) Examples of TEM figures. B) The angular distributions obtained from TEM results. C) Double peaks fitting of the angular distribution. The lower peak is $32.7^{o}$ and the higher peak is $62.7^{o}$.

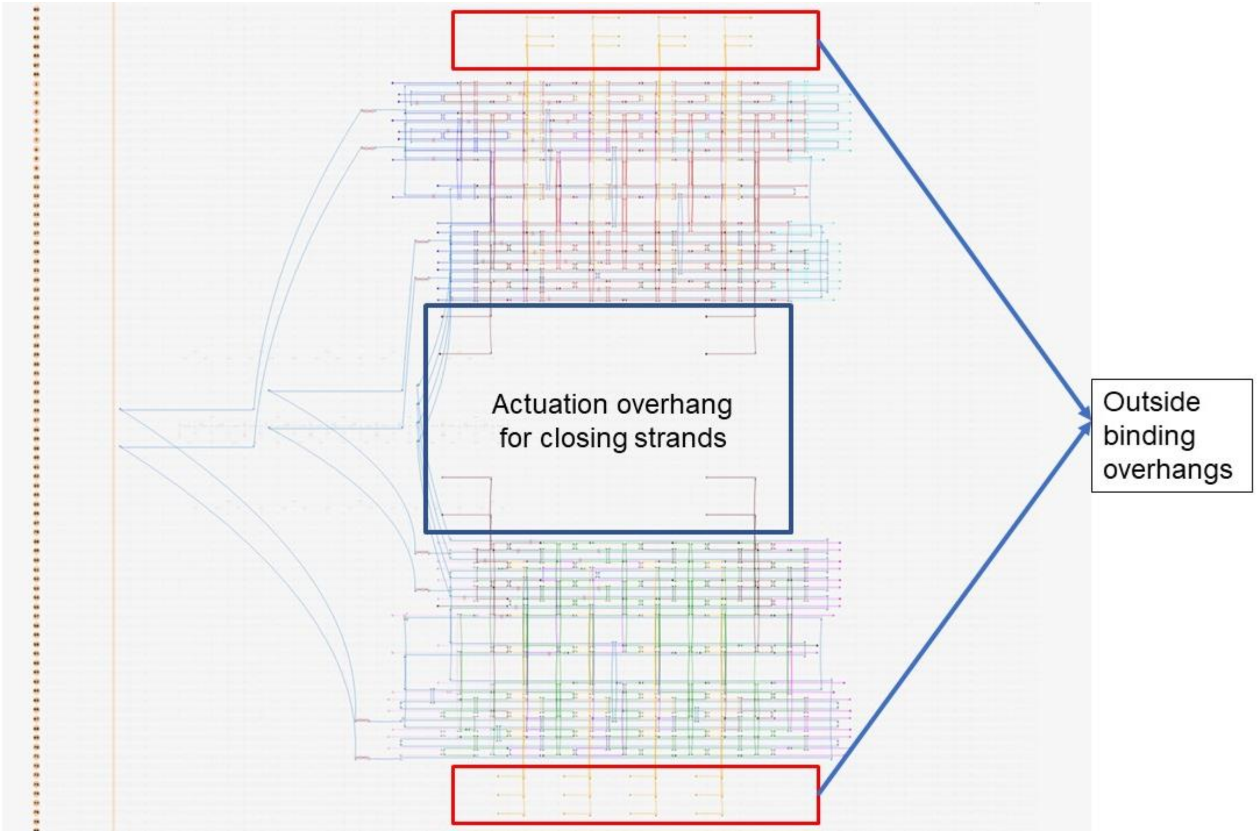

**Figure S6.** The caDNAno schematics of DNA origami hinge for self-assembling.

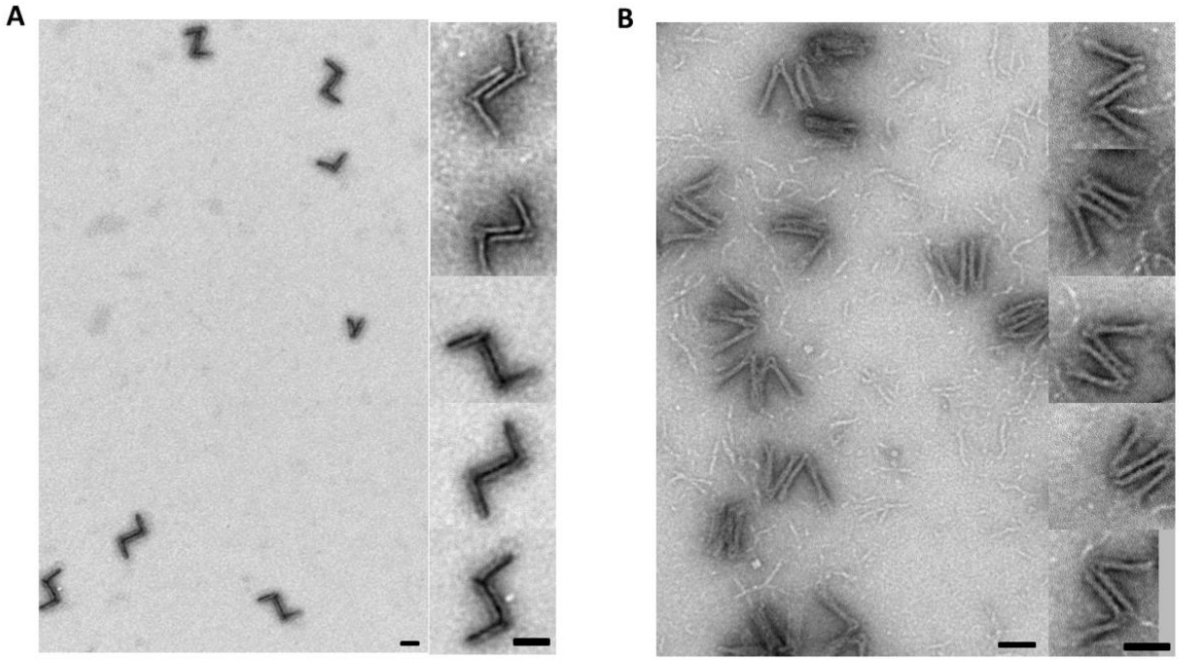

**Figure S7.** Examples of TEM figure for dimers of DNA origami hinge linked by coiled-coil peptides. A) The overhangs are set on the inner side of the arm. B) The overhangs are set on the outer side of the arm. There are two different assemble directions when two hinge binding together.

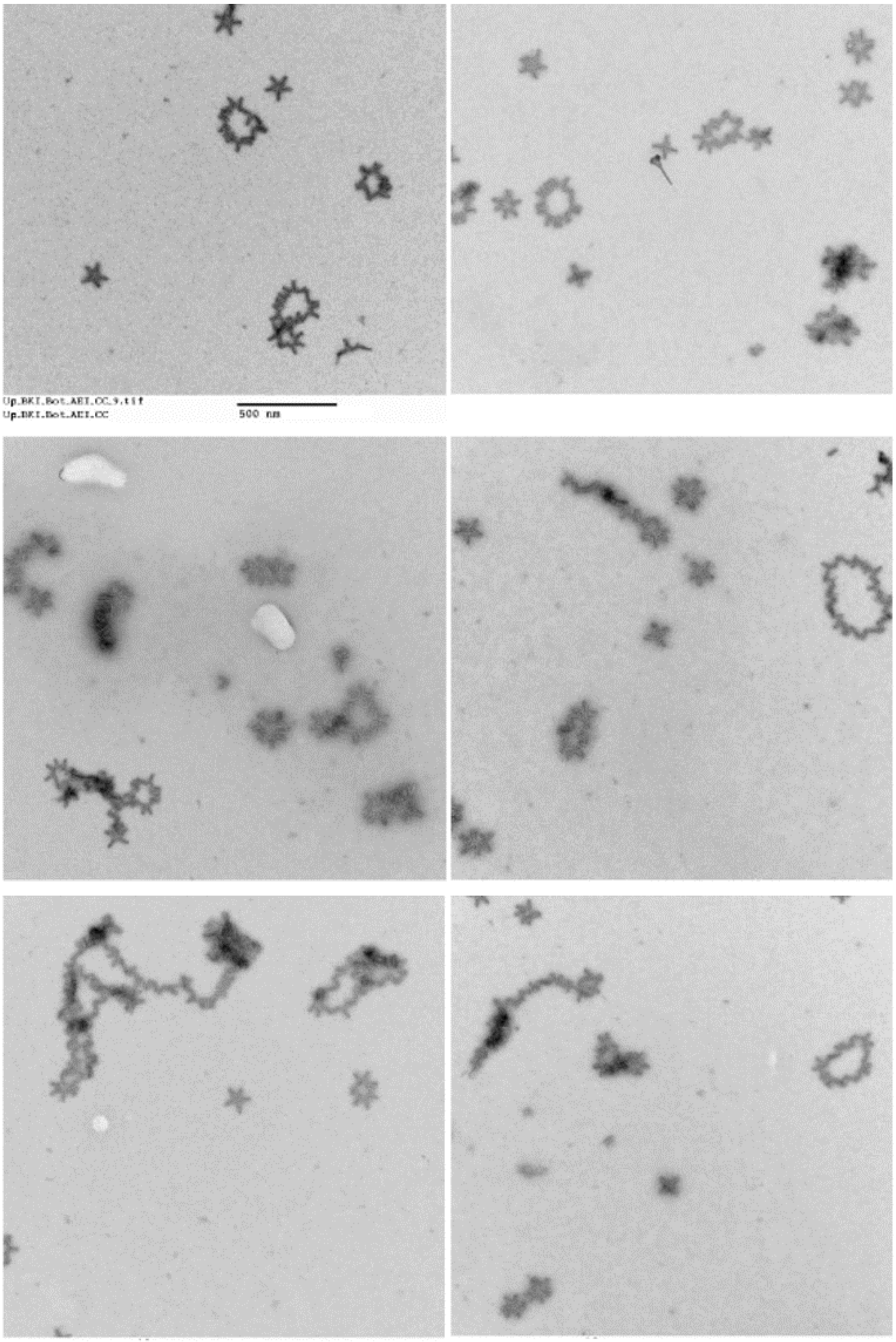

**Figure S8.** Representative TEM images of unrestricted self-assembly with coiled-coil peptides. Both polymers and circle patterns can be observed.

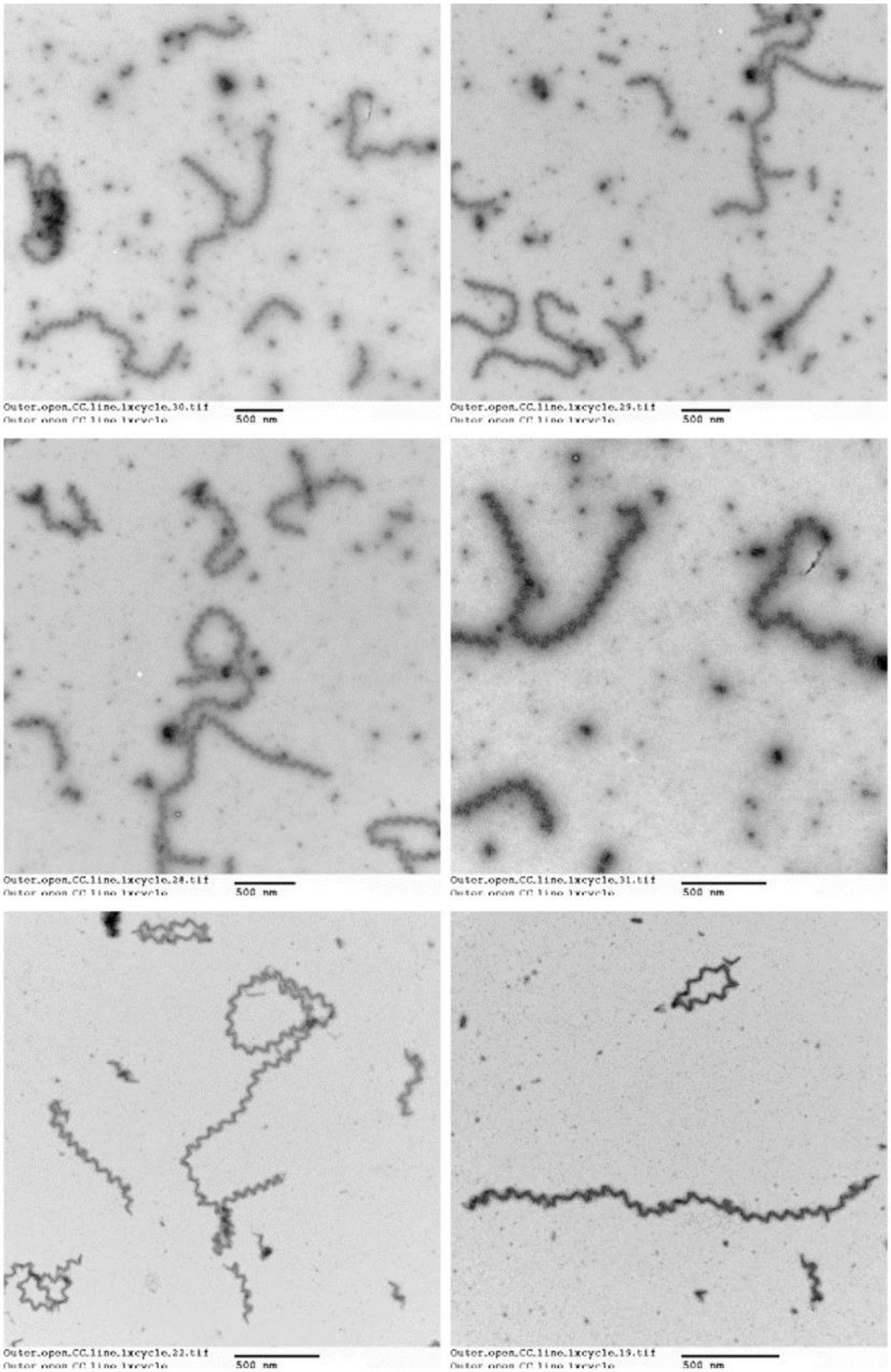

**Figure S9.** Representative TEM images of linear polymers self-assembly with coiled-coil peptides.

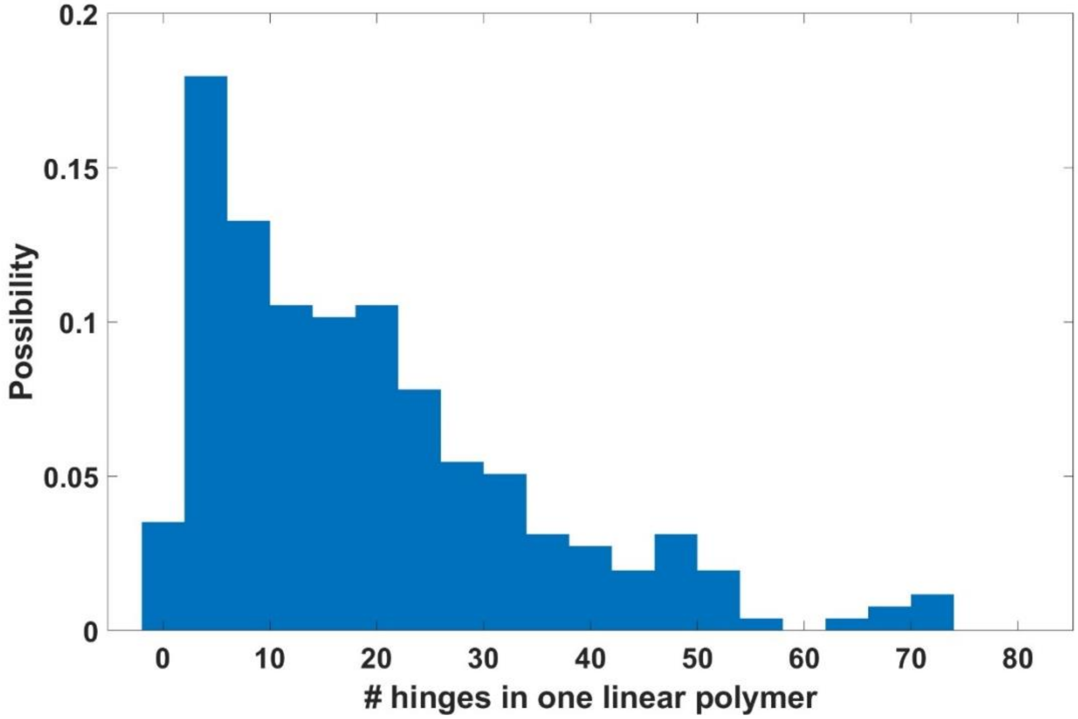

**Figure S10.** Length analysis of linear polymers in TEM images.

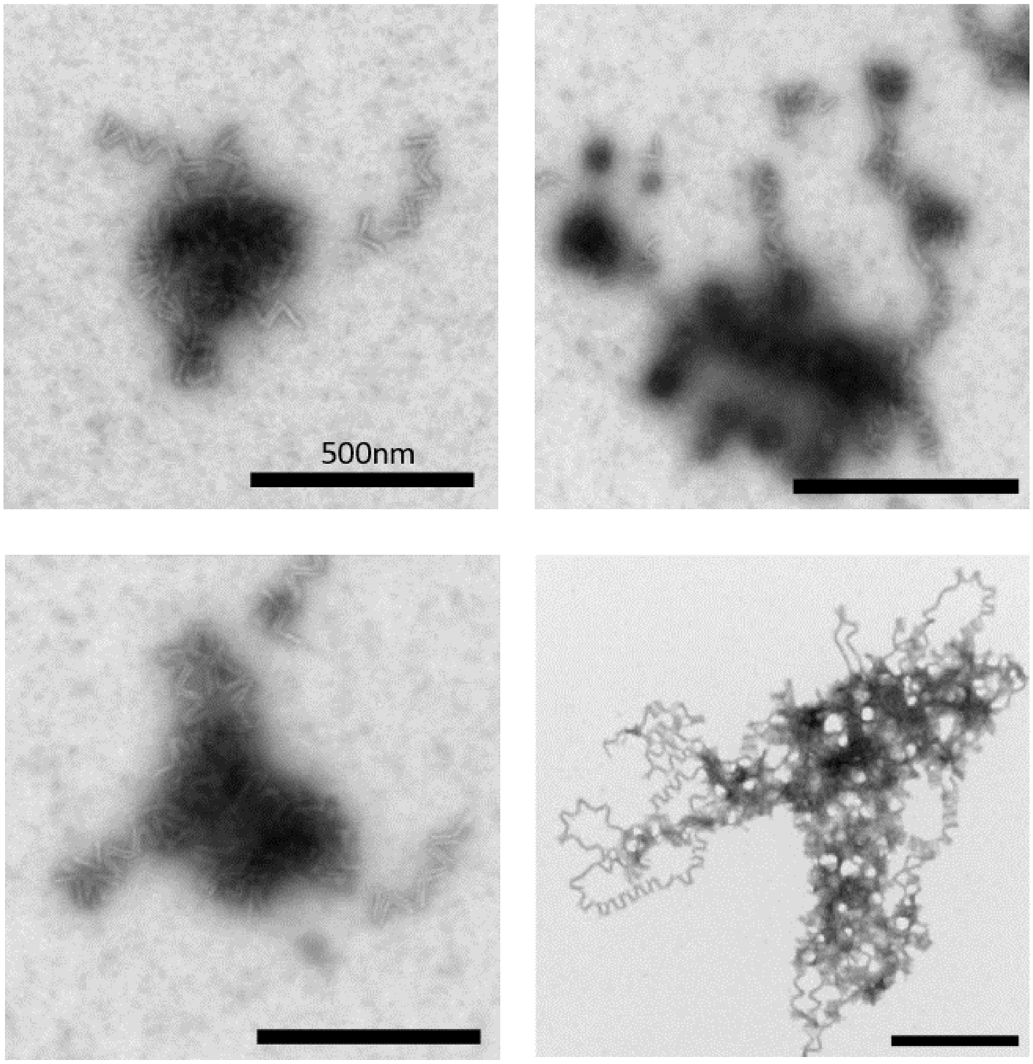

**Figure S11.** Representative TEM images of linear polymers self-assembly with replacement DNA strands. In linear polymer self-assembly with coiled-coil peptides, 20x access AEI and BKI were added to 5nM origami structure, and in the comparison experiment only with replacement DNA strands, 20x access of single strand DNA was added to 5nM structure.

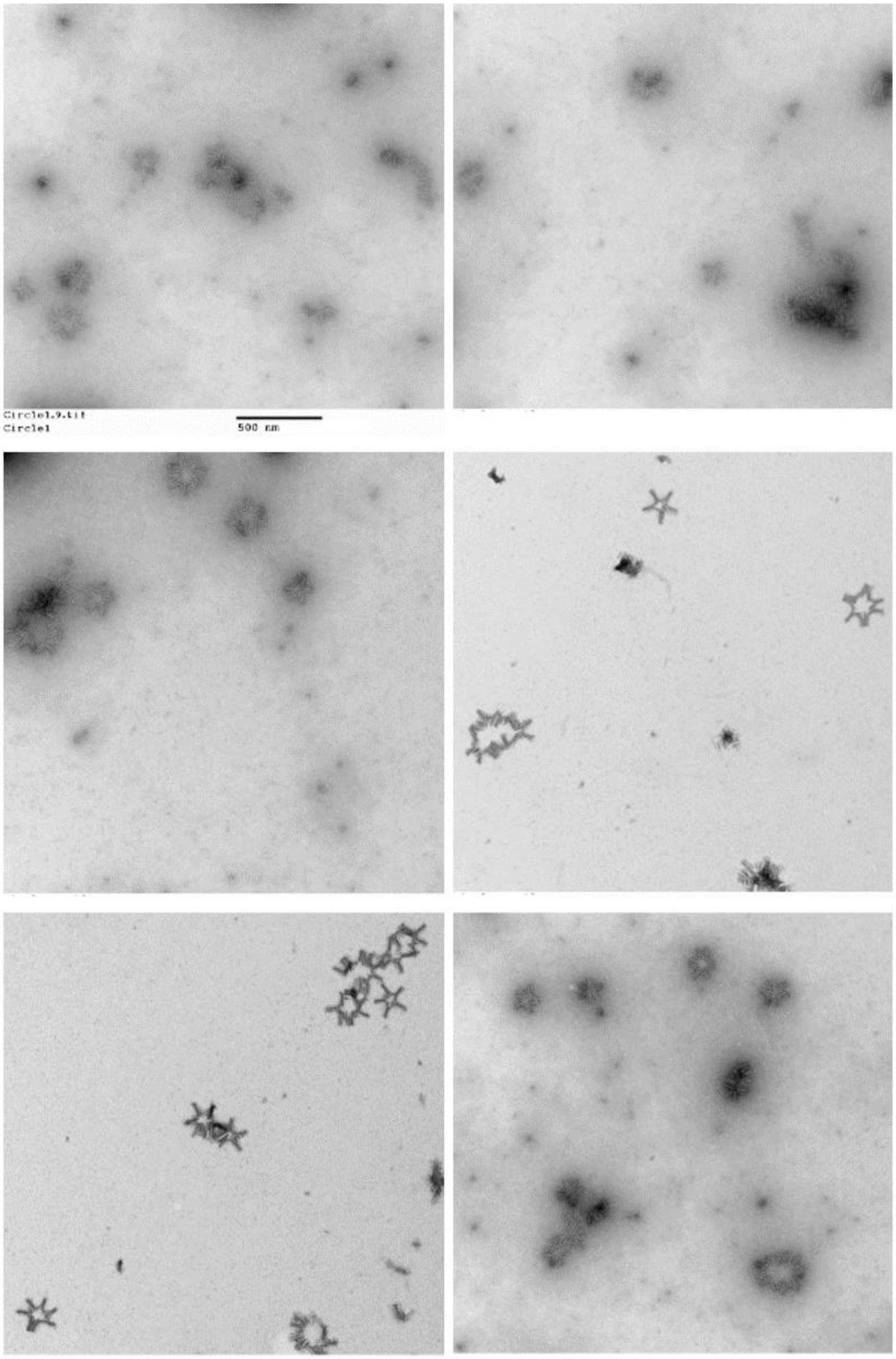

**Figure S12.** Representative TEM images of failed circle pattern self-assembly with coiled-coil peptides. It does not form clear circle pattern as design and both same and opposite direction assemblies happen.

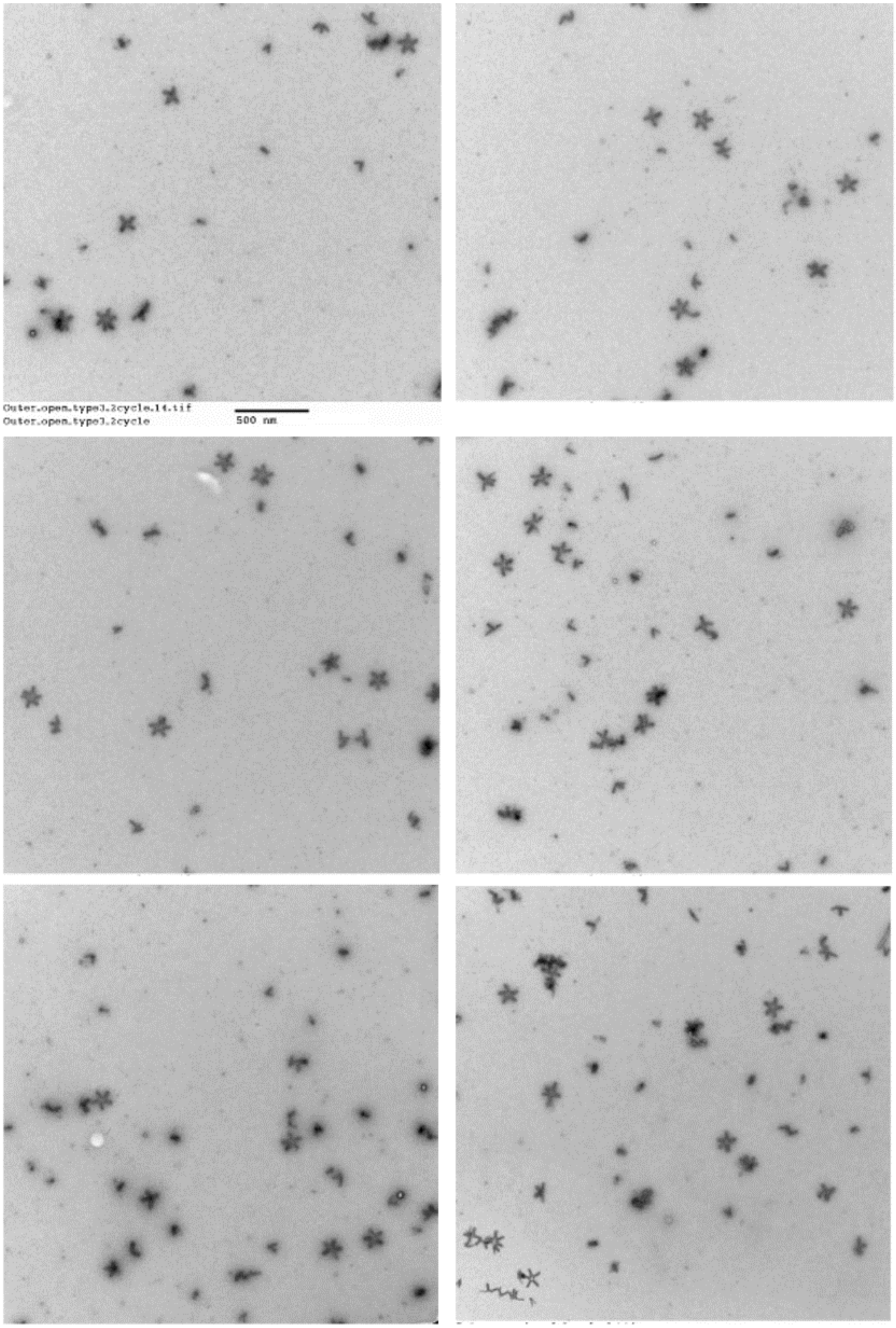

**Figure S13.** Representative TEM images of complexed overhangs design of circle pattern self-assembly with coiled-coil peptides. Clear circle patterns can be observed.

**A** **B**

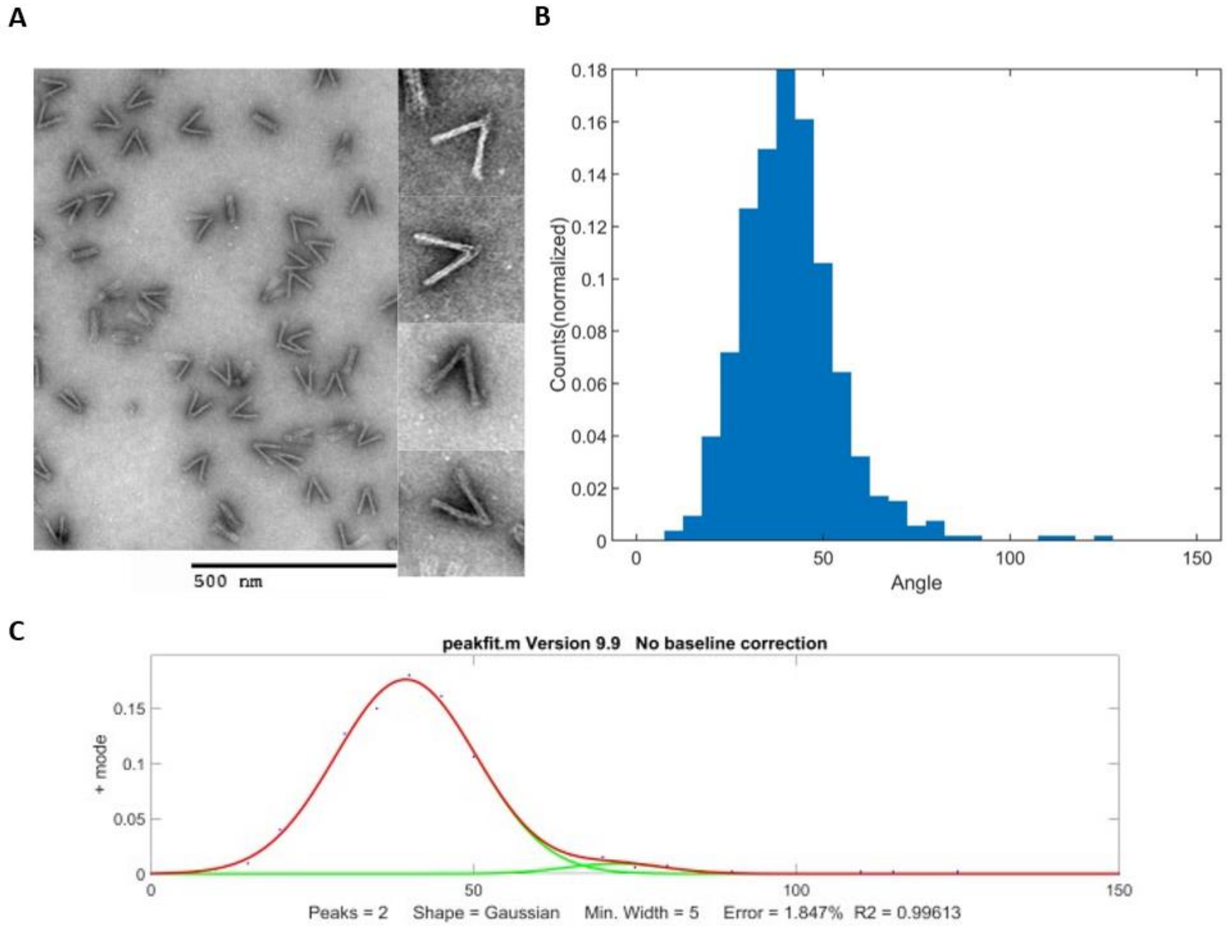

| | Position |
|---|---|
| High peak | 71.9 |
| Low peak | 39.5 |

**Figure S14.** Examples TEM figures of actuated hinge with average angle 41$^{o}$ before self-assembly. B) The angular distributions obtained from TEM results. C) Double peaks fit the angular distribution. The lower peak is 45$^{o}$.

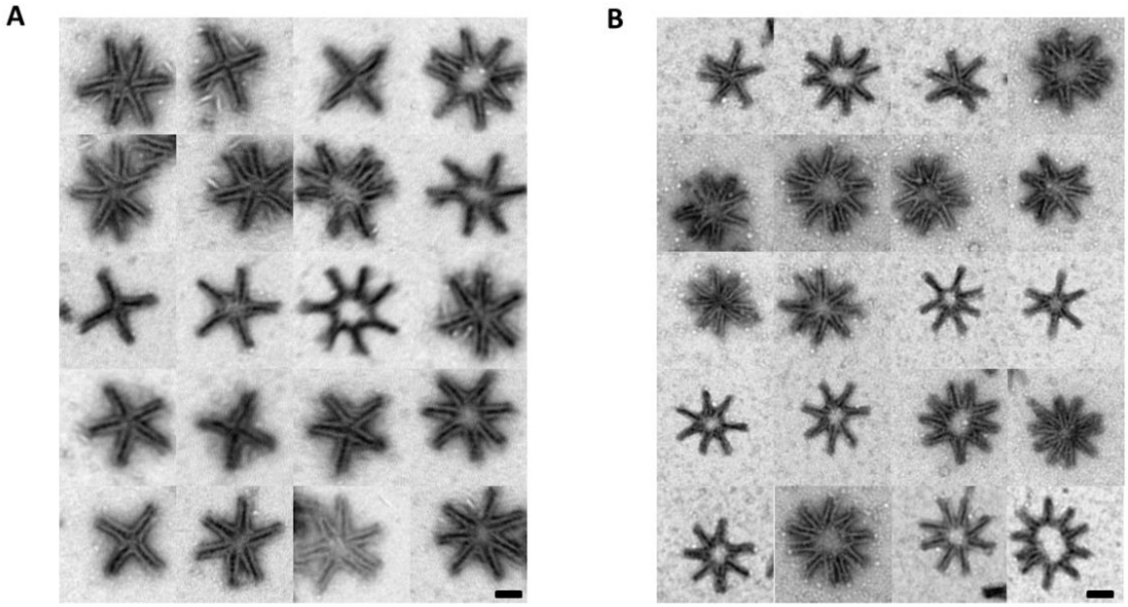

**Figure S15.** Examples TEM figures of self-assembly circular pattern with actuated hinge A) Circle pattern formed by non-actuated hinge self-assemble. B) Before self-assembling, the hinges are actuated to 45$^{o}$. Then obtain the circle pattern through self-assembly.

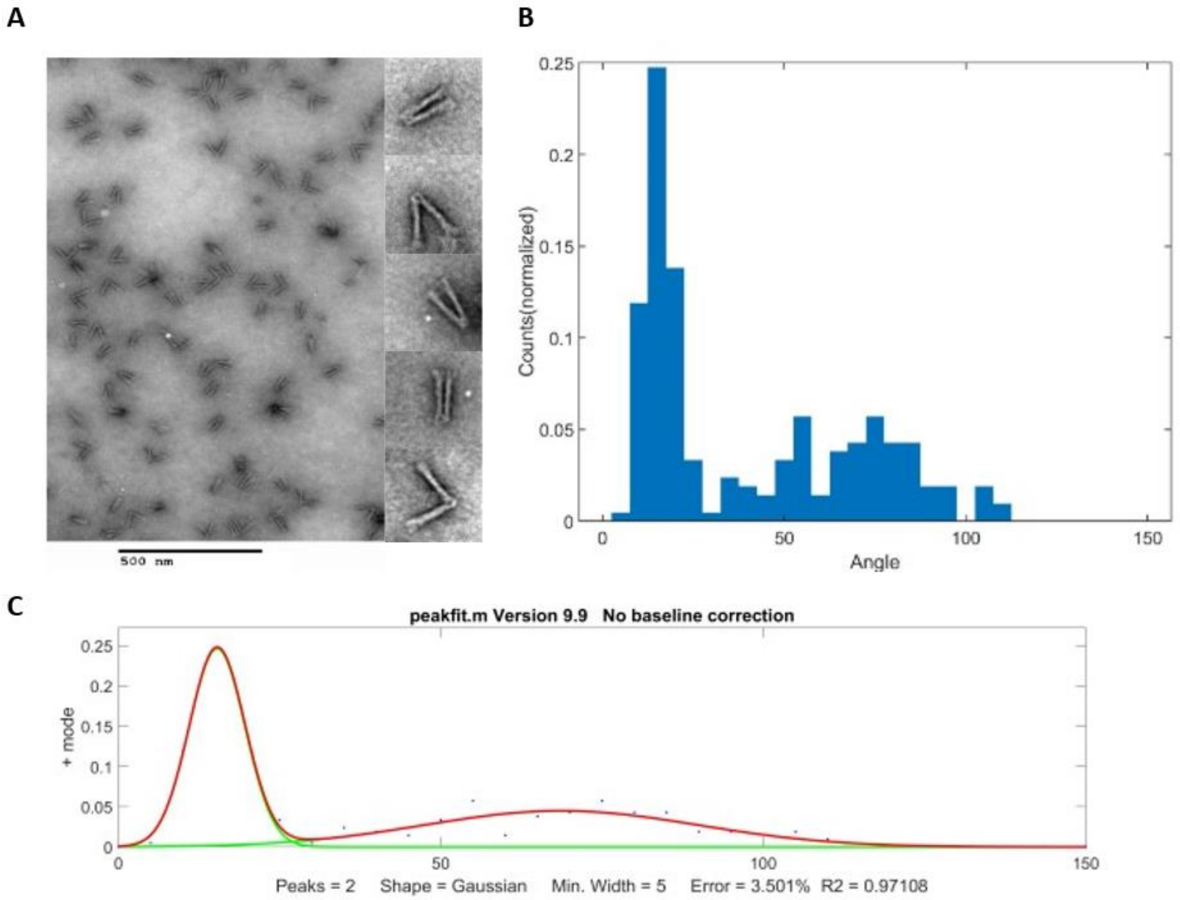

| | Position |
|---|---|
| High peak | 68.67 |
| Low peak | 15.32 |

**Figure S16.** Examples TEM figures of actuated hinge which is used in self-assemble linear polymer. B) The angular distributions obtained from TEM results. C) Double peaks fit the angular distribution. The lower peak is 15$^{o}$.

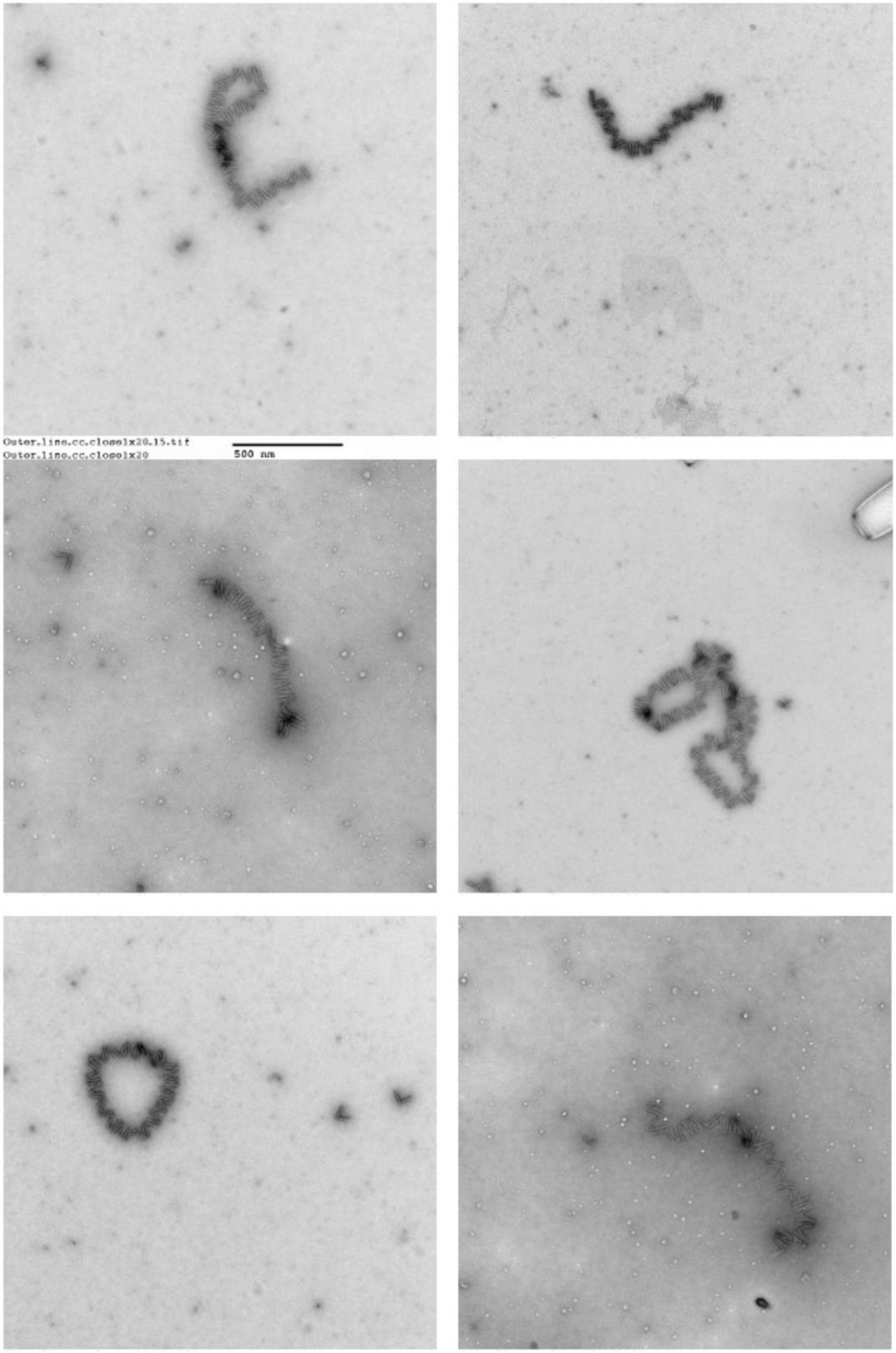

**Figure S17.** Examples TEM figures of actuated linear polymers.

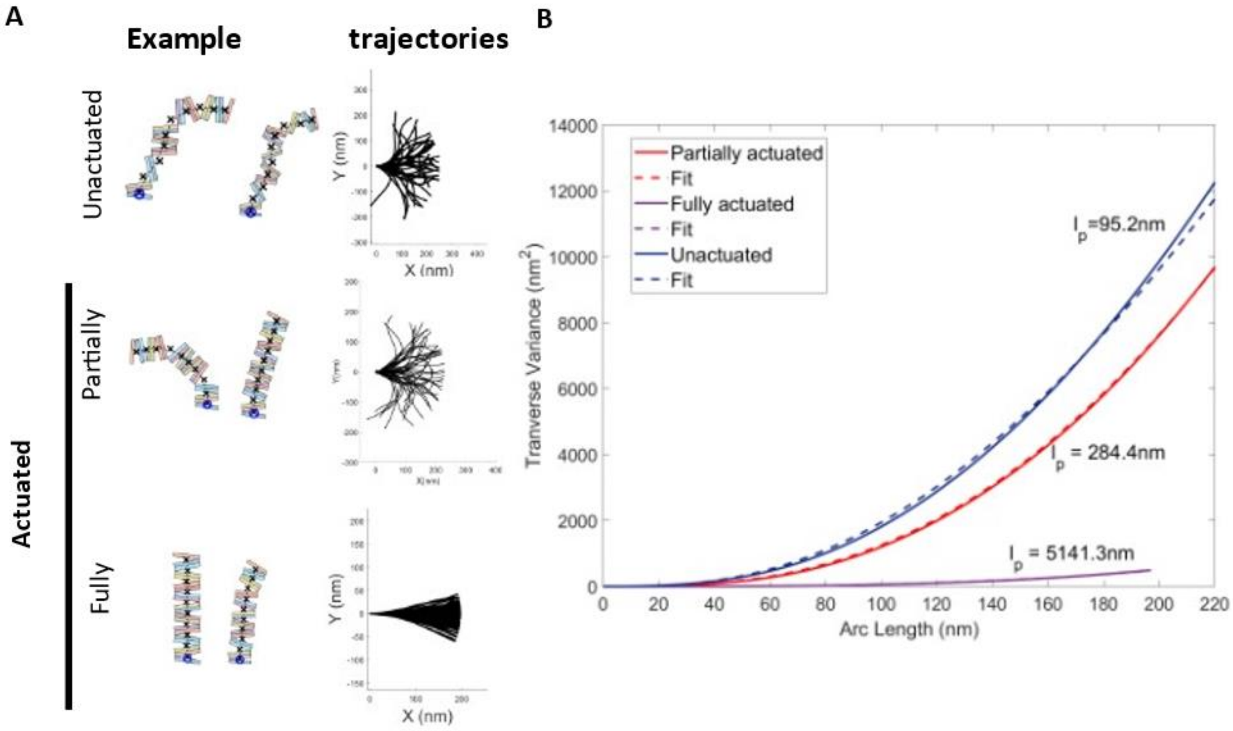

**Figure S18.** Persistence length measurements of simulations of unactuated, partially actuated and fully actuated linear polymers. A) Examples of 10 hinges assembled polymers with unactuated and actuated hinge. Partially actuated polymer means the simulation was built on the angle distribution of actuated polymers and fully actuated polymer means the simulation was built on the angle only picked up from angle less than 30 degrees in the same distribution. B) The transverse fluctuations were fit to equation (1) to give a persistence length.

| Sequence | Element name |
|---|---|
| CGGCGAACGTGGCGAGTTCTTTTCACCAGTGAGGGAGAGG | Arm staple 1 |
| AAGGAACAACCACAGACAATATTTCATTGTAGC | Arm staple 2 |
| AGAAACAAGGTAATTGAGCGCTAACCTTTACA | Arm staple 2 |
| TCTGTCCAGGCCGATTAAAGGGATGACTCCAACGTCAAAG | Arm staple 3 |
| AACCGGATATTCAAATCGCGAAACAAAGTACATCGTCACC | Arm staple 3 |
| TTTTTTTTAAGTTGGGTTGTGCACTCTGTTTTTT | Arm staple 4 |
| TATGTAAACGAACAAATTCATTAAAGGTGAATTTAGAGCC | Arm staple 4 |
| CGCCTCCCCCCCCTTATTAGCGTTAGCAAAGCGGATTGCA | Arm staple 5 |
| TATCGGCCCGGATTCTCCGTGGGAGTAGGTAA | Arm staple 5 |
| GAGAGAATAAGCCGTTTTTATTTTCATCGTAGATAATCGGCTGTCTTTCGGTCAT A | Arm staple 6 |
| TTTTTTCAGGGTGGTTTAAAGGAAGCTGGCAAGTGTAGCGGCAGGAAAA | Arm staple 6 |
| TTTTTTACACAACATACGAGGCTGGAGGTGTC | Arm staple 7 |
| TTTTTTTGGTGCTGCGGCCAGAATGCGGCGGGCAGTGTCAC | Arm staple 7 |
| TAATGGGTCTGTGTGATAAACAA | Arm staple 8 |
| TGAAATAGCAATAGCTCAGATAGCTGCTGATGTTTTTT | Arm staple 8 |
| AGCATCGGCAGCGATTATACCAAGAGCGAGAGGCTTTTGCAAAAGAAGCGTT TACC | Arm staple 9 |
| TTTTTTTACGTAATGCCACTACATTAAACGGGTAAAATTTTTT | Arm staple 9 |
| AGCCGCACGAACGTGCCGGACTTGCCTGTCGTGGCCCTGA | Arm staple 10 |
| GTAATCGTATCAGTTGACACTATCATAACCCTTTTT | Arm staple 10 |
| TTAAATGCCTTTATTTCAACGCAAGGATAAAAATTTTTTT | Arm staple 11 |
| CCTGAGTATTGCTTTGACGAGCACGGTCGAGGTGCCGTAA | Arm staple 11 |
| TTTTTTCCAGAACCACCACCAGAGCCGCCGCCATCAGAGCCACCGGAAC | Arm staple 12 |
| ACCCGCATTGACAGGAGGTTCAAACAAATTTTTAATGGAA | Arm staple 12 |
| CATTGCAATCACGCTGCGCGTAACGGGAAAGC | Arm staple 13 |
| ACGCTCATGAAATGGATTATTTACTCGAGGTG | Arm staple 13 |
| CCACGGAATAGCGCGTTTTCATCGGAAGCCCGAAAGACTT | Arm staple 14 |
| TGTACATCTGCTGGTCTGGTCAGCTTGCGCTCCTGGTTTG | Arm staple 14 |
| CACCAGAAGTGTACAGTGCTCCATGTTACTTACCACGCAT | Arm staple 15 |
| AGAATCCTTTGGCCTTTTTAACCAATAGGAAGAGGGTAG | Arm staple 15 |
| TTTTTTAGCGGGCGCTAGGGCGGGAAGAAAGCGAAAGGTTTTTT | Arm staple 16 |
| AGACGACGCATTATTACAGGTAGAAAGATTCGTCAGTGC | Arm staple 16 |
| ATAAGAGGAAAGTACGGTGTCTGGCTGTAATACTTTTGCG | Arm staple 17 |
| AAAGTTAAGCAGCCTC | Arm staple 17 |
| AGAGGTGAATTCACCACCGCCAGC | Arm staple 18 |
| TAATTGCGAGCAACCGCAAGAATTTGCCGCCAGCAGTTGTCGA | Arm staple 18 |
| TTCGCAAAGCGTTTTACTTTAGCGTCAGACTGTAAGTTTA | Arm staple 19 |
| CTGGTAATCTTAATGCGCCGCTACCCTAAAGGGAGCCCCC | Arm staple 19 |
| CGGAACCTGAAGAAAAAGCTGCTCATTCAGTG | Arm staple 20 |
| GATTTAGATTGCCCTTCACCGCCTGCCAGCTGCATTAATG | Arm staple 20 |

| | |
|---|---|
| AACCGATAGTTTATCAGCTTGCTTATTGGCAGGGCGGTCAGTATTAACACCTTTTTT | Arm staple 21 |
| GGATTAGAAACAGTTGATTCCCAAGCTAAATCGGTTGTAC | Arm staple 21 |
| TTTTTTTGACGCTCAATCGTCTGGAAATACCTACATTTTTTTTT | Arm staple 22 |
| TTTTTTGAGGAAGGTTATTCCGGCAAACTTTTTT | Arm staple 22 |
| ATGCAACTTCATTTTTCACCAGTAGCACCATTATATTGAC | Arm staple 23 |
| TTTTTTTTAGTTAACGAATTCACAAACCGGAATCATTTTTTGTT | Arm staple 23 |
| TACCGAGCTACACTGGTGTGTTCAACAATCGGCGAAACTGCT | Arm staple 24 |
| TTTTGCACATAAGAGAATATAATTTTTT | Arm staple 24 |
| GGCTGACCAATAAGGCTTGCCCTGTCATTATA | Arm staple 25 |
| TTTTTTGGCATCAATTCTACTAATAGTAGTAGCAT | Arm staple 25 |
| CATTAAATTTTTGCTATCAGGTCATTTTTTT | Arm staple 26 |
| CTGCTCATGCCAACGGCAGCACCGCCTAATGAGATGGTG | Arm staple 26 |
| TCAAAAAGTGGGGCGCGAGCTGAAAAGGTTTTTTT | Arm staple 27 |
| GTCGCTATTCGCGTCTGGCCTTCCAACCGTTCTAGCTGAGAATTAGAGATACAT | Arm staple 27 |
| TCAAATCAAGCATAAATTCTGCGA | Arm staple 28 |
| CGAACTAAACAGTTAATGCCCCCTCCCTCAGAACCGCCAC | Arm staple 28 |
| AACCCTCATAGCCCGGAATAGGTGAAAGTATT | Arm staple 29 |
| TTTTTTTTTAGAACCCTCACGTTGGTGTTTTTTT | Arm staple 29 |
| TAGAAGCAAAAGAAGTTACATACCAGTATAAAGCCAGCCTAATT | Arm staple 30 |
| ATGGCTATTAGTCTTTTCGTATTAGATGATAC | Arm staple 30 |
| TTTTTTAGATGGGCGCACTGCAAGGCGATTTTTT | Arm staple 31 |
| CTTGAGTAAATAAGTTTTAACGGGGTCATACAACAACGCCTTATTTGA | Arm staple 31 |
| GTGCATCTCCGTAATGGGATAGGTCATATATT | Arm staple 32 |
| CGGATTGAGCCAGTTTTATAACTAACAAAGAA | Arm staple 32 |
| TTTTTTTTCAACTTTAATCATTCTTGAGATGGTTTAATTTTTTT | Arm staple 33 |
| CAGTAAGCAAAACTAGCATGTCAAGATGAACG | Arm staple 33 |
| TTTTTTATCACCGGAACCAGAGCCACCACCACCCTCAGAGCCGCCATTTTTT | Arm staple 34 |
| AATTTCTTTGACAACAACCATCGCGCCGGAACGAGGCGCAACTTTGAA | Arm staple 34 |
| TAACGTCAACCGCGCCCAATAGCAAGCAAATCATCCTAATTTACGAGCGCCTGTTC | Arm staple 35 |
| TACTCAGGGGACTTGCTGAACCTCATAGATAATACATTTGGTAAAAAA | Arm staple 35 |
| TTCGCGTCCGTGAGCCCATCAGATGCCGGGTTGAGCCGCC | Arm staple 36 |
| CCGGGGGTTTCTGCCAGCCGGTGCCCCCTGCAAAACGACG | Arm staple 36 |
| AGCAAAATGCGGATGGCTTAGAGCTTAATTGCTGAATTTTTTT | Arm staple 37 |
| TTTTTTCAGTTTGGAACAAGAGTCCACTATTAA | Arm staple 37 |
| GGTCATAGCTGTTTCAAAGGTTTCTTTGCTACCAGTCC | Arm staple 38 |
| CAAAAGGGAACCATCGATAGCAGCAAACTCCAACAGGTCA | Arm staple 38 |
| AATACTTCTCGTTAGAATCAGAGCTCATGAACCATCACCC | Arm staple 39 |
| TTTTTTAGTACCAGGCGTGAAAGGAATTTTTTTT | Arm staple 39 |
| TAGTTTGACCATTCAAAATTAAGCAATAAAGCCTCAGCCATCAAT | Arm staple 40 |
| TTTTTTCAAATCCAGGGATGTGTCGTAACC | Arm staple 40 |

| | |
|---|---|
| CTCAGCAGAATAATTTTTCACGTCCCTTCTGAGCCCTAAAACATCGCGAAGTATTCCACCCTC | Arm staple 41 |
| TGAGAGACATCGCACTTGTTGGGAAGGGCGATCAAGCTTT | Arm staple 41 |
| ATACGTGGTAAAGGAAGTCACCAG | Arm staple 42 |
| TTTTGTCATAGAAAATACATACATTGAATACCAAGATGATGAAACAAATCAATATA | Arm staple 42 |
| AAGAGGCTGAACTGGCACGAGAAA | Arm staple 43 |
| TTCCAAGAGAACAAGCAACATAAAAACAGGGACTGAACAAAGTCAGAG | Arm staple 43 |
| CCATGTACCAGAGCCAAGACTTTACAAACAATGGCGGTTG | Arm staple 44 |
| CCTCAAGTGTACTGGTACAGTGCCCGTATAACGGAACAAATAAAAAC | Arm staple 44 |
| TTTTTTCATTTTCGAGCCAGTACCAGCTACAATTT | Arm staple 45 |
| TCAGATGAATATACAGTAACAACATGTAATTTAGGCAGAGGTTTTTT | Arm staple 45 |
| AAGTTTCCGAAGGCACCAACCTAAGCGTCCAATACTGCGGAATCGTCATAAATATTCA | Arm staple 46 |
| TGCCAGTTGTTTTAGCGAACCTCCCGACTTGCGCTAATGCAGAACGCGGTAATCAT | Arm staple 46 |
| TTTTTTATAATGCTGTAGCTCAACATGTTTTAAAT | Arm staple 47 |
| AGCACTAACAGCAAGCGGTCCACGACTGCCCGCTTTCCAG | Arm staple 47 |
| GCCCGAGATAGGGTTGAGTGTTGTTCTTTTTT | Arm staple 48 |
| CGGTTTGCGTATTGGGCGCTTTTTT | Arm staple 48 |
| TTTTTTCCTGAGAGTCTGGTAATGCAGATACTTTTTT | Arm staple 49 |
| TTTTTTTTTTAAGAAAAGTAAGATCTTACCGAAGCCCTTTTTTT | Arm staple 49 |
| TTAAATCAAGATTAGTGTCCAGAC | Arm staple 50 |
| ACGAGTAGCCGGAAGCACCGTAAT | Arm staple 50 |
| TATCCTGACATATTTAACAACGCCAGTACCTTTTACATCGCGCC | Arm staple 51 |
| TTTTTTGATAGCTCTCACGATCATTTTGCGGATTTTTT | Arm staple 51 |
| TTGCGGGAACGGAGATTTGTATCAGAGTAATCTTGACAAG | Arm staple 52 |
| AGGGAACCGCCACCCTCGTAACACAACTGATACCTGAAAATCACTTG | Arm staple 52 |
| CTCAGAACGGAGAAACTTAATTACATTTAACAATTTCTTTTTT | Arm staple 53 |
| CAGAGGCGAGTATGTTCTGAGAAGAGTCAATACTGGTGCCGGAAAATC | Arm staple 53 |
| TTTTTTGCGGTCCGTTTTTTCGTCTCGTCGCTGACGATGCT | Arm staple 54 |
| GATTGCTTAAAGGTGGTGAAAACATAGCGATTTTCCCTTTGTGAGTGAATAACCTGTACAGCG | Arm staple 54 |
| CAAATATCTGGTCAATAACCTGTTCAATAAATCATACAGG | Arm staple 55 |
| CAGCCCTCTCTGAATTAGTTTGAGTAACATTGAAAAAGAGACGCAGAACGCAAC | Arm staple 55 |
| TCACCGGAAGCAAATCGTTAACGGTCCTCACAAGAAAAAT | Arm staple 56 |
| CCAGCCAGCTTTCCGGTCAACATTAAATGTGAGGAGACAG | Arm staple 56 |
| AGATTCAATTATGACCAAGTTTCA | Arm staple 57 |
| TTCATCAATCGCCTGATAAATTGTCTTGCAGG | Arm staple 57 |
| CCAGTCAGGACGTTGGATTATTCTGAAACATGTATCACCG | Arm staple 58 |
| GCTCATTTGATATTCAGAGGCAGGAAACAAAAAATAACGGCTTAATTG | Arm staple 58 |
| CTATTTTTTAACATCTAGCTATATTTTCATTATTAAGAG | Arm staple 59 |

| | |
|---|---|
| CAAAAACAAAGGGTGAGAAAGGCCGCGAGTAACAACCCGTTCAGGAAGTACC | Arm staple 59 |
| ATTTTCAGAGGTTTAGTACCGCCAGCCTATTT | Arm staple 60 |
| CCTTATGCGATTTTAAGAGACTCCTCAAGAGAAGGGTTGA | Arm staple 60 |
| ACCCTGACTATTATAGTCAGATGCCATCTTTTCATAATCAAATTTTTT | Arm staple 61 |
| AGAGGACAGATGAACGCGAGTAGTAAATTGGGGTGAATTA | Arm staple 61 |
| ATGATATTCTGTAGCCAGCTTTCACACCGCTTGTGAATTTAATGGTTT | Arm staple 62 |
| TATACTTCGTATGGGATCTAAAGTTTTGTCGTCTTTCCAGACTTTTTT | Arm staple 62 |
| CAATTACCCAAATCAACGTAACAAATCTACGTTAATAAAA | Arm staple 63 |
| TGGGGTGTCGGTGGTGCCATCCCAACAGCGG | Arm staple 63 |
| AACTTTTTGCCTCTTCGACGACAG | Arm staple 64 |
| TTTTTTGGAACCGAACTGACCAGACGGTCAATCATAAGTTTTTT | Arm staple 64 |
| TATCATATTATTATTTATCCCAATAAGGCTTA | Arm staple 65 |
| GCGTTAAAAGAACTGGCATAGGTC | Arm staple 65 |
| TTTTTTGCCTGCAACAGTGCCACGCCTGGTCAGGAGCACTAACAACTAATGAAGGGT | Arm staple 66 |
| GGAAATTAGTTACCAGAAGGAAACTAAGAGCACGCGAGAA | Arm staple 66 |
| GAGTTAAAAAAGGCTCCAAAAGGATAAAAGGGCGAACCACCAGCAGAAAGCCGTCAAAATATCA | Arm staple 67 |
| CCCAATAACGAGGAAAGGCTTAGGTTGGGTTAGAGGGGACGCTATTAC | Arm staple 67 |
| TCCAGCGCCGTTTTCACCTTATCA | Arm staple 68 |
| GAAATTATTCATTTCATAACCAGGCAAAGCGCCATTCGCCGGATAACC | Arm staple 68 |
| TTTTTTGCCAGAATGGAAATGATAATCAGAAA | Arm staple 69 |
| GCATTTTCGGTCATAGTCAGAGCCGCCAAACGAAAAGACC | Arm staple 69 |
| TTTTTTATAACGCCAAAAGGAATTACGGACTGGATAAACGAAAGAGGCAAAAGGACTAA | Arm staple 70 |
| CAGTAGCGCATATGGTTTACCAGCAGACTCCTTATTACGCAATACCGACCGTGTGACTGTTTAG | Arm staple 70 |
| CCATGTTTCGTCATAAACATCCCTTCGAATTCCCTGTTTA | Arm staple 71 |
| TTTTTTGTTAGTAAATGAATTTTCTTGAATAATGGAAGGGTTAG | Arm staple 71 |
| GACGACAAAATTGTTATCCGCTCACAATTTTTTTT | Arm staple 72 |
| GATTGCCGTCTAAAATATCTTTAGTTGGCAAA | Arm staple 72 |
| TTTTTTTTGAGCCATTTGGGAATATCACCGTCACCGACTTTTTT | Arm staple 73 |
| AGAATCGCATCTTACCAACGCTAATTGAAGCC | Arm staple 73 |
| TACAAACTTGGCTTTTAATCCTTTGCCCGAATTAAATTT | Arm staple 74 |
| CAGAGGTGACCTGCAGCCAGCGGTGCACGCGTATGTAGAA | Arm staple 74 |
| TCACTGTTGCCCTTCTCCGTGGTGAATTTTTT | Arm staple 75 |
| CAGAAGGATCCTGATTTAAAAGAG | Arm staple 75 |
| TGCGCGCCAACGCCAGGGTTTTCCGCGAAAGGATCGCAAG | Arm staple 76 |
| GAAGGTAAACCATTAGCAAGGCCGTTAATTGCTCCTTTTG | Arm staple 76 |
| AATCGGCCGCACATCCTCATAACGAGGCGGCC | Arm staple 77 |
| AGAAAATTACAGAATCAAGTTTGCATTCGAGCTTCAAAGCGAACCAGAATT | Arm staple 77 |

| | |
|---|---|
| GCCAACTCATCTTTGACCCCAACGAGGGTAGCAACATAGA | Arm staple 78 |
| GGCGAAAAATCGGCAAAATCCCTTCATAAAGTGTAAAGCC | Arm staple 78 |
| AGATAGAATGAAAATCATAGGAAC | Arm staple 79 |
| ACCAGTAAGCCTTTAAATGAAAAA | Arm staple 79 |
| TCTAAAGCATCACTAGATACCGAAACATTCTGCGGCCTTG | Arm staple 80 |
| TTTTTTATAGTTGCGCCGACAAAAACAGCTTGATACCGTTTTTT | Arm staple 80 |
| TTCCATATGAGTACCTGAAACGTCACCAATGACGACATTC | Arm staple 81 |
| AGGCTTTGAGAATACACTAAAACGAGGGGGT | Arm staple 81 |
| GAATACCACATTCAACAGCAAACAAGAGAATCTCATATGT | Arm staple 82 |
| TTTTTTATTTGAATTACCTTAAATCCTCATTATTTTTT | Arm staple 82 |
| TTTTTTACAAAGAAACCACCGTAACGATTTTGCTTCATGAGG | Arm staple 83 |
| TCAACAGTGATAAGTGCCGTCGAGAGGATTAGGATTAGCGGGGTTTTGCTCTT TTTT | Arm staple 83 |
| TTTTTTAAGTGTTTTTATAATACGCCAGAATCCTGAGTTTTTT | Arm staple 84 |
| AAATCAAGGGCGAAAATCCTGTTTGTGAGCTAACTCACAT | Arm staple 84 |
| CGCTGAGGGTCGAAATCCGCGACCACCAGGCGCATAGGCT | Arm staple 85 |
| TTTTTTAGTACCGACAAAAGGTAAAGTAATTCTTGCTA | Arm staple 85 |
| TTTTTTGTACCGCACTCATCGAACGGGTATTAAACCAATTTTTT | Arm staple 86 |
| CAAGGCAAATAAATTAATGCCGGACGCCATCAAAAATAATTAATTAATAGCT | Arm staple 86 |
| AAAGAATAAGAACGTGTTTAGACAGGAACGGTCAGTGAGGCCACCGAGGTTT GGAT | Arm staple 87 |
| CAATTCATTTTCAGCGTCCACAGA | Arm staple 87 |
| AATAGTAAAATGTTTAAGGCATAGTAAGAGCAAGATTTAG | Arm staple 88 |
| TTTTTTGAATTAACTGAACACCAGCGCATTAGACGGGATTTTTT | Arm staple 88 |
| TCGGGAAATAGAACGTCAGCGTGGGACATAAA | Arm staple 89 |
| AGACTTTTAAACAACTTTCAACAGCAATATAAGCGGAATTATCATCATATTTTAA ATACCGTTC | Arm staple 89 |
| AACCGATTTAACGGAATACCCAAATAAGAATAAATTTCATCTTCTGACTGCGCA AC | Arm staple 90 |
| TCCGGTATTCTAAGAAAGATAAGTCCTGAACAGTTGAGGATCCCCGGG | Arm staple 90 |
| GTTCCGAAACCGTCTAGGGAGCTATTT ATA TGG TCA ACT G | uparm_1_1_BKI |
| AACAGGATCACGCAAATTAACCGGATGATGGTTT ATA TGG TCA ACT G | uparm_1_2_BKI |
| ATCAAACCGTTATTAATTCCTGATGTAGCATGAGTGAGAGGCTACAGTTT ATA TGG TCA ACT G | uparm_1_3_BKI |
| CCCCAGCATTTTTTGGGTATAACGTTT ATA TGG TCA ACT G | uparm_2_1_BKI |
| TGCTTTCCTTTGATTAGTAATAACGCGTAAGATTT ATA TGG TCA ACT G | uparm_2_2_BKI |
| CAACAATGCGCGTGAGTTTCTTGCGAATCGAAAGACTTT ATA TGG TCA ACT G | uparm_2_3_BKI |
| GAGAGTTGATCGGAACAGGGCGCGTTT ATA TGG TCA ACT G | uparm_3_1_BKI |
| TACTATGGGAAGAACTCAAACTATGCCAACAGTTT ATA TGG TCA ACT G | uparm_3_2_BKI |
| AAAATCCCAGGATTTACATTAAAACAAGCCCATCCAAAAAGGCCGCTTTTT ATA TGG TCA ACT G | uparm_3_3_BKI |

| | |
|---|---|
| ACAGCTGAGCTTGACGCACCACACTTT ATA TGG TCA ACT G | uparm_4_1_BKI |
| CCGCCGCGATCCAGAACAATATTAGTCACACGTTT ATA TGG TCA ACT G | uparm_4_2_BKI |
| TTTAGTGATAGATTAGGATAAAACAGCAGCAATTGTATCGTATTCGGTTTT ATA TGG TCA ACT G | uparm_4_3_BKI |
| GTTCCGAAACCGTCTAGGGAGCTATTGT AAT ACC AGA TGG | uparm_1_1_AEI |
| AACAGGATCACGCAAATTAACCGGATGATGGTTGT AAT ACC AGA TGG | uparm_1_2_AEI |
| ATCAAACCGTTATTAATTCCTGATGTAGCATGAGTGAGAGGCTACAGTTGT AAT ACC AGA TGG | uparm_1_3_AEI |
| CCCCAGCATTTTTTGGGTATAACGTTGT AAT ACC AGA TGG | uparm_2_1_AEI |
| TGCTTTCCTTTGATTAGTAATAACGCGTAAGATTGT AAT ACC AGA TGG | uparm_2_2_AEI |
| CAACAATGCGCGTGAGTTTCTTGCGAATCGAAAGACTTGT AAT ACC AGA TGG | uparm_2_3_AEI |
| GAGAGTTGATCGGAACAGGGCGCGTTGT AAT ACC AGA TGG | uparm_3_1_AEI |
| TACTATGGGAAGAACTCAAACTATGCCAACAGTTGT AAT ACC AGA TGG | uparm_3_2_AEI |
| AAAATCCCAGGATTTACATTAAAACAAGCCCATCCAAAAAGGCCGCTTTTGT AAT ACC AGA TGG | uparm_3_3_AEI |
| ACAGCTGAGCTTGACGCACCACACTTGT AAT ACC AGA TGG | uparm_4_1_AEI |
| CCGCCGCGATCCAGAACAATATTAGTCACACGTTGT AAT ACC AGA TGG | uparm_4_2_AEI |
| TTTAGTGATAGATTAGGATAAAACAGCAGCAATTGTATCGTATTCGGTTTGT AAT ACC AGA TGG | uparm_4_3_AEI |
| CATGTTCAGGGAGGTTCGAGCGTCTTT ATA TGG TCA ACT G | bottom_1_1_BKI |
| TTTCCAGAACGCTCAACAGTAGGGATTCGCCTTTT ATA TGG TCA ACT G | bottom_1_2_BKI |
| CGGAATTTTACATAAACATCAAGATCAGACGACAACATATCAAAGACATTT ATA TGG TCA ACT G | bottom_1_3_BKI |
| TCAACAATCGCGAGGCACAAAATATTT ATA TGG TCA ACT G | bottom_2_1_BKI |
| AACAGCCAGCGTTATACAAATTCTAAATCGCGTTT ATA TGG TCA ACT G | bottom_2_2_BKI |
| TCTGATTACCTGTTAAGACGAGCAAACGCAATCAATTTT ATA TGG TCA ACT G | bottom_2_3_BKI |
| AATATCCCAGATATAGCCAAATAATTT ATA TGG TCA ACT G | bottom_3_1_BKI |
| GAAACGATAATTACTAGAAAAAGCTAAATAAGTTT ATA TGG TCA ACT G | bottom_3_2_BKI |
| ACGGGAACATTCAGGCCTAAATTTATCAAAATCATGATTAGCCAAAGATTT ATA TGG TCA ACT G | bottom_3_3_BKI |
| ACCAATCAGAATCATTAAAATGAATTT ATA TGG TCA ACT G | bottom_4_1_BKI |
| AATAGCAGTATCAGAGAGATAACCGAGTTAAGTTT ATA TGG TCA ACT G | bottom_4_2_BKI |
| GCCAGTGCCGGTGCGGCAAATATATAACCTCCCGCAATAAGAGGGAGGTTT ATA TGG TCA ACT G | bottom_4_3_BKI |
| CATGTTCAGGGAGGTTCGAGCGTCTTGT AAT ACC AGA TGG | bottom_1_1_AEI |
| TTTCCAGAACGCTCAACAGTAGGGATTCGCCTTTGT AAT ACC AGA TGG | bottom_1_2_AEI |
| CGGAATTTTACATAAACATCAAGATCAGACGACAACATATCAAAGACATTGT AAT ACC AGA TGG | bottom_1_3_AEI |
| TCAACAATCGCGAGGCACAAAATATTGT AAT ACC AGA TGG | bottom_2_1_AEI |
| AACAGCCAGCGTTATACAAATTCTAAATCGCGTTGT AAT ACC AGA TGG | bottom_2_2_AEI |
| TCTGATTACCTGTTAAGACGAGCAAACGCAATCAATTTGT AAT ACC AGA TGG | bottom_2_3_AEI |
| AATATCCCAGATATAGCCAAATAATTGT AAT ACC AGA TGG | bottom_3_1_AEI |

| | |
|---|---|
| GAAACGATAATTACTAGAAAAAGCTAAATAAGTTGT AAT ACC AGA TGG | bottom_3_2_AEI |
| ACGGGAACATTCAGGCCTAAATTTATCAAAATCATGATTAGCCAAAGATTGT AAT ACC AGA TGG | bottom_3_3_AEI |
| ACCAATCAGAATCATTAAAATGAATTGT AAT ACC AGA TGG | bottom_4_1_AEI |
| AATAGCAGTATCAGAGAGATAACCGAGTTAAGTTGT AAT ACC AGA TGG | bottom_4_2_AEI |
| GCCAGTGCCGGTGCGGCAAATATATAACCTCCCGCAATAAGAGGGAGGTTGT AAT ACC AGA TGG | bottom_4_3_AEI |
| GGTCATAGCTGTTTCAAAGGTTTCTTTGCTACCAGTCCTTGT AAT ACC AGA TGG | Group 1 bottom1 |
| AGAATCCTTTGGCCTTTTTAACCAATAGGAAGAGGGTAGTTGT AAT ACC AGA TGG | Group 1 bottom2 |
| TGGGGTGTCGGTGGTGCCATCCCAACAGCGGTTT ATA TGG TCA ACT G | Group 1 up1 |
| TGGGGTGTCGGTGGTGCCATCCCAACAGCGGTTT ATA TGG TCA ACT G | Group 1 up2 |
| CCGGGGGTTTCTGCCAGCCGGTGCCCCCTGCAAAACGACGTTGT AAT ACC AGA TGG | Group 2 bottom1 |
| TATCGGCCCGGATTCTCCGTGGGAGTAGGTAATTGT AAT ACC AGA TGG | Group 2 bottom2 |
| AATCGGCCGCACATCCTCATAACGAGGCGGCCTTT ATA TGG TCA ACT G | Group 2 up1 |
| AACCCTCATAGCCCGGAATAGGTGAAAGTATTTTT ATA TGG TCA ACT G | Group 2 up2 |
| AAGACCCACTCAGCTTACGCCGGAAGATAAATCA | Circle_actuate_overhang_up1 |
| AAGACCCACTACCCCGGTGCGCAGTCATAGTTAG | Circle_actuate_overhang_up2 |
| ACAGGTGAGAGATAGACTTGCGGCTGGCGTGCGTGTG | Circle_actuate_overhang_bot1 |
| GAGAGATCTACAAAGGTTAAATCACGTGCGTGTG | Circle_actuate_overhang_bot2 |
| AGT GGA CCA GTG GGT CTT TTC ACA CGC ACG | Circle_closing strand |
| AGTCGGGTCTATACGAAGACCCACTCGGCCAGAAACGCGCGGACGGGCA | Polymer_actuate_overhang_up1 |
| AGTCGGGTCTATACGAAGACCCACTTATAAGTAATCAATATTGAGAGCC | Polymer_actuate_overhang_up2 |
| GGAGAAGCAATGCCTGAGTAATGTACAAACGGCGTGCGTGTGGACCCACTTAGGCCC | Polymer_actuate_overhang_bot1 |
| GCCAGCTGCAGTCACGACGTTGTATCAGACGACGTGCGTGTGGACCCACTTAGGCCC | Polymer_actuate_overhang_bot2 |
| AGT GGA CCA GTG GGT CTT CGT ATA GAC CCG ACT TTG GGC CTA AGT GGG TCC ACA CGC ACG | Polymer_closingstrand |

Table S1 Sequence information of DNA origami structure.
